# Tumors accumulate expanded GATA3-dependent tissue Tregs

**DOI:** 10.64898/2026.03.13.707380

**Authors:** Katharina Kunesch, Sraddha S. Bharadwaj, Jacqueline L.E. Tearle, Lydia Kopplin, Isaac Desveaux, Asmae Laouina, Fabio Ticconi, Andreas B. Wild, Makoto Mark Taketo, Marc P. Stemmler, Thorsten Cramer, Gesine Hansen, Ulf Neumann, Kylie R. James, Oliver Pabst, Ana Izcue

## Abstract

Targeting Tregs is a potential strategy to improve cancer therapies. However, which Tregs accumulate in response to tumoral processes, and how tumors affect their phenotype, is poorly understood. Here we show that tumor Tregs are equivalent to effector tissue Tregs in steady state organs. We used a mouse model of intestinal neoplasia to demonstrate that one early event in carcinogenesis is sufficient to induce local accumulation of Tregs resembling human tumor Tregs. Treg accumulation was driven by TCR-dependent oligoclonal expansion of tissue Tregs with an effector Treg phenotype. Treg expansion was independent of CCR8, IL33R and CD137, which were previously linked to tumor Treg. In contrast, GATA3 was required for effector tissue Tregs and for their expansion in response to neoplasia. Our findings identify GATA3-dependent clonal expansion of effector tissue Tregs as a key event in promoting tumor growth.

**Highlights:**

- An early tumorigenic event alone drives accumulation of effector tissue Tregs
- Tregs in tumors are phenotypically akin to effector tissue Tregs
- The accumulation of Tregs is driven by TCR-dependent oligoclonal expansion
- GATA3 controls tumor-promoting effector tissue Tregs

## Introduction

Tumors develop in close connection with the local immune microenvironment, in a crosstalk that shapes both compartments. Immune cells exert context-dependent effects on tumors that can both promote and suppress tumor growth. CD4^+^ Foxp3-expressing regulatory T cells (Tregs) are key to establishing and maintaining immune tolerance to tumors ^1^. Indeed, high levels of Foxp3^+^ Tregs are generally associated with bad prognosis in human solid tumors and, in mouse models, accumulation of Tregs actively favors tumor growth and prevents immune rejection of the tumor ^1^. Despite the high potential of Treg manipulation in cancer therapy, how Tregs react to early tumoral changes and which subsets respond to them is still broadly unknown.

Since the identification of Foxp3 as the transcription factor driving Tregs, it has become evident that Tregs are heterogeneous in terms of functional capacities, requirements and location preferences^2^. A large proportion of Tregs in blood and lymphoid organs belong to the lymphoid type expressing CCR7, high levels of CD62L and low CD44. Lymphoid Tregs typically express the transcription factor Helios and their maintenance is independent of TCR/antigenic signals but dependent on cytokines ^3,4^. Upon activation, Tregs undergo transcriptional changes that enhance migration into tissues ^5^. Tissue Tregs maintain Helios expression and can further differentiate, acquiring transcriptional profiles often associated to the expression of conventional T-helper transcription factors such as T-bet and GATA3. Tissue Tregs often adopt a transcriptional profile specific to their tissue of residence, and tissue Treg populations can show diverse immunosuppressive activity ^6,7^. Among activated Tregs, tissue Tregs expressing the inhibitory receptor KLRG1 show a distinctive phenotype and high immunosuppressive capacity. We term these cells effector tissue Tregs. In addition to these subsets, some tissues, such as the gut, harbor a population of microbiota-dependent, peripherally induced Helios^-^ RORγt^+^ Tregs.

Intrinsic Treg heterogeneity is a confounding factor in the study of tumor Tregs. Indeed, while most studies acknowledge that the phenotype of tumor Tregs differs from Tregs in other organs, there is not a consensus “tumor Treg signature”. This is partly due to differences in Treg phenotype across different tumors, but also to differences in the organ chosen to isolate “control Tregs”. In addition, single cell analysis, often used in this setting, does not always contain enough Tregs to allow side-by side comparison of Treg subtypes in control and tumor tissue, and prioritizes the discovery of highly expressed genes, which can lead to important transcription factors, which typically have lower expression, being overlooked. Finally, the analysis of the interplay between nascent tumors and Tregs is complicated by the low number of Tregs in the early stages of tumorigenesis and the need to precisely time tumor development. The general consensus is that tumor Tregs present an activated phenotype, but it is unclear whether they are unique, or if they correspond to a subset that is present in normal tissue. If so, it would be important to identify their counterpart in healthy tissue, and understand how the tumor environment affects their activation.

Here we study Treg responses to neoplastic changes using a mouse model of inducible neoplasia. We find that a single tumorigenic event, which mimics the initial tumorigenic event in many cases of colorectal cancer ^8^, drives the local accumulation of Tregs akin to human tumor Tregs. Strikingly, this Treg accumulation was characterized by extensive TCR-dependent oligoclonal expansion of effector tissue Tregs with no signs of additional immune activation compared to their steady-state counterparts. We found that effector tissue Treg response to neoplasia is independent of receptors associated with tumor Tregs, such as IL-33R, but effector tissue Tregs rely on GATA3 during steady state. Importantly, biallelic GATA3 non-redundantly controls tumor-promoting effector tissue Tregs.

Collectively, our data show that the initial oncogenic events already trigger the GATA3-dependent local accumulation of effector tissue Treg clones with tumor-supporting ability, which may set the basis for the immunosuppressive tumor microenvironment.

## Results

### Mouse intestinal neoplasia induces the accumulation of effector tissue Tregs resembling human tumor Tregs

To study changes in the immune environment and Treg populations during early stages of tumorigenesis, we used the mouse model of tamoxifen-induced stabilization of β-catenin in intestinal epithelial cells ^8^. This model recapitulates early molecular pathways in colorectal cancer development that affect the entire gut and that we refer to as gut neoplasia (Figure 1A).

**Figure 1.**
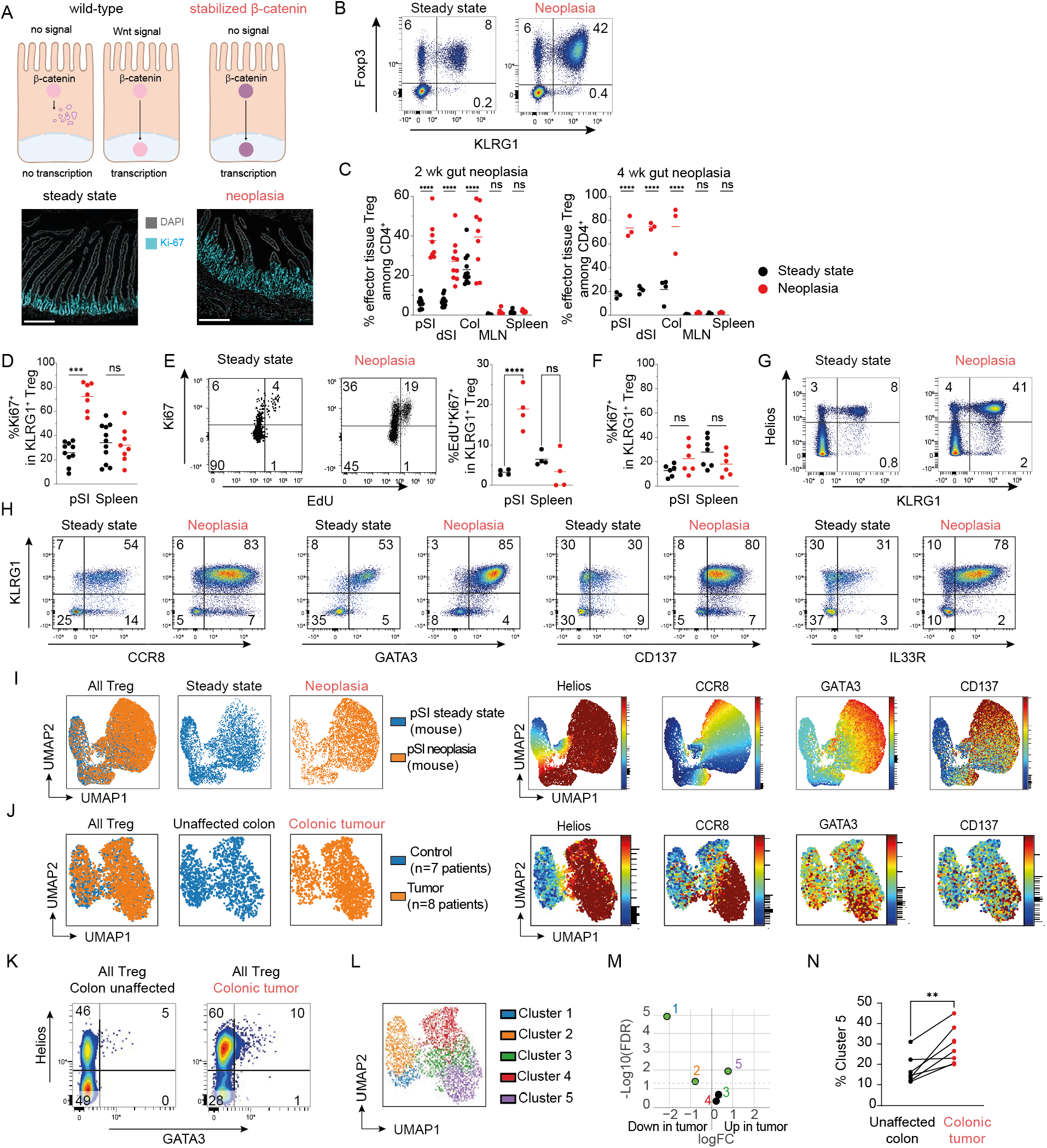
Mouse intestinal neoplasia induces the accumulation of Treg resembling human tumor Treg. (A-I) Analysis of mouse model of gut neoplasia. (A) Diagram and microscopy images depicting neoplastic changes induction in the intestinal epithelium by tamoxifen treatment in the *Catnb*^*+/lox(ex3)*^ Vil Cre-ERT model. Tamoxifen injection leads to the expression of β-catenin lacking exon3, which is a stabilized form not targeted for degradation. Microscopy shows DAPI and Ki67 staining of proximal small intestine of control mice and *Catnb*^*+/lox(ex3)*^ Vil Cre-ERT mice 2 weeks after tamoxifen injection. Bar: 200 µm. (B-C) FACS plots (B) and frequencies (C) of effector tissue KLRG1^+^ Tregs in control mice (black symbols) and mice with tamoxifen-induced intestinal epithelial neoplastic changes (red symbols) at 2 and 4 weeks after neoplasia induction. Gated on CD4^+^ live T cells from proximal small intestine (gating strategy in Figure S1B). FACS plots show changes at 2 weeks post neoplasia induction. (D Frequencies of Ki67^+^ cells among effector tissue KLRG1^+^ and control KLRG1^-^ Tregs at 2 weeks after neoplasia induction (E) Representative FACS plots and frequencies of proliferating EdU^+^ Ki67^+^ cells from controls and mice with gut neoplasia 2 weeks after neoplasia induction. FACS plots show proximal small intestine. (F) Frequencies of Ki67^+^ cells among effector tissue KLRG1^+^ Tregs at 4 weeks after neoplasia induction (G) Representative stainings of Helios versus KLRG1 among CD4^+^ T cells of the proximal small intestine 2 weeks after induction of neoplasia. (H) Representative stainings of CCR8, IL-33R, CD137 and GATA3 versus KLRG1 among Foxp3^+^ Tregs of the proximal small intestine 2 weeks after induction of neoplasia. (I) UMAP of the Foxp3^+^ Tregs in (H). UMAP was generated on KLRG1, CCR8, Helios, CD137 and GATA3 (J-N) FACS analysis of Treg from human colonic tumors. Gating strategy in Figure S1F. (J) UMAP analysis of concatenated Tregs from matched tumoral (orange, 8 samples) and unaffected tissue (blue, 7 samples). UMAP was generated based on expression of CCR8, CD137, PD-1, CD25, Foxp3, GATA-3, CD161, CTLA-4, CD127 and HELIOS. (K) FACS staining showing expression of HELIOS and GATA3 in intestinal Tregs (L) Analysis showing clustering of the concatenated Treg in (J) (M) differences in cluster frequencies between Tregs from tumors or from unaffected tissue (N) frequencies of cluster 5 among Tregs from the indicated organs. **, p<0.01, unpaired t test.

Characteristic of the neoplastic gut was a massive disturbance of gut architecture, with extensive expansion of crypts and increased number of proliferating epithelial cells. These changes were progressive with milder phenotype at 2 weeks (Figure 1A) and stronger alterations of gut architecture at 4 weeks (Figure S1A). The changes took place along the whole intestinal tract, from proximal small intestine to colon, and were accompanied by a striking increase in Tregs in the intestinal lamina propria. In accordance with related models of intestinal neoplasia ^9^, this increase was driven by effector tissue Tregs identified as Foxp3^+^ KLRG1^+^ CD4^+^ T cells (Figure 1B, C). A related model with increased epithelial Wnt signaling showed that neoplasia induces effector tissue Tregs independently of microbiota (Figure S1C). Accordingly, we observed the Treg increase all along the intestinal tract, from the microbiota poor proximal small intestine to the microbially rich colon (Figure 1C). Neoplasia progression further increased effector Treg frequencies to make up the majority of the intestinal CD4^+^ T cell compartment 4 weeks after tamoxifen injection (Figure 1C). In contrast, the frequency of KLRG1^-^ Treg subsets remained unaffected (Figure S1D). The accumulation of effector tissue Tregs correlated with subset-specific increase in proliferation, as assessed by Ki-67 expression (Figure 1D). Furthermore, incorporation of EdU was most pronounced in effector tissue Tregs (Figure 1E), suggesting that indeed local expansion rather than cell recruitment caused changes in Treg numbers and composition. Strikingly, the increase in proliferation was transient and returned to basal levels at 4 weeks post tamoxifen injection (Figure 1F) despite high effector Treg accumulation.

Neoplasia-responsive Tregs showed a phenotype associated with “tumor Tregs”. They expressed Helios and GATA3, as expected for KLRG1^+^ Treg in the gut (Figure 1G). They also expressed the canonical markers of tumor Treg signature CCR8, IL-33R (ST2) and CD137 (4-1BB) ^10-14^. Indeed, the observed increase in these markers among Tregs upon neoplasia was mostly driven by the expansion of KLRG1^+^ effector tissue Tregs, rather than by their general upregulation in the Treg population (Figure 1H, S1E). These data suggest that the markers associated with “tumor-Treg signature” are not specific for tumors, but correspond to the phenotype of effector tissue Tregs in the gut.

We then used a high-parameter flow cytometry panel to check if the effector tissue Treg phenotype we observe in the mouse model (Figure 1I) is also found in Treg accumulating in human colonic tumors. There are some fundamental differences between mouse and human gut Tregs. In general, Treg frequencies are lower in human gut as compared to mouse intestine ^15^, and KLRG1 cannot be used as an effector Treg marker since it is not expressed by human Tregs (^16^ and Figure S1F-G). We therefore used a panel including Foxp3, Helios, CCR8, CD137 and GATA3 to compare the phenotype of accumulating Tregs from colorectal tumors and from unaffected colonic tissue from the same patients (Figure 1J). We did identify a GATA3^+^ Treg population in human colon, although at much smaller frequency compared to the mouse gut (Figure 1K). Cluster analysis identified 5 Treg clusters among human colonic Treg cells (Figure 1L). In general, tumors harbored lower frequencies of Helios negative Tregs (clusters 1 and 2) (Figure 1M, S1H), which are putatively peripherally-induced microbiota-dependent Tregs. In contrast, the Helios^+^ GATA3^+^ Treg cluster 5, enriched for markers expressed by mouse effector tissue Tregs, was expanded in colonic tumor samples as compared to unaffected tissue (Figure 1M-N), suggesting that tumor-induced accumulation of effector tissue Tregs is a conserved feature between mice and humans.

### Single cell analysis of intestinal Tregs fails to identify a tumor-induced signature on neoplasia-responding Tregs

To further define the changes in effector tissue Treg during early gut neoplasia, we extended the analysis to single cell RNA (scRNA-seq) and TCR sequencing (scTCR-seq). Single donor mice were used to enable the analysis of TCR-clones at high resolution in individual mice. To account for the massive differences in Treg numbers between steady state and neoplastic intestines, effector tissue Tregs were enriched as CD4^+^ CD25^+^ KLRG1^+^ cells. CD4^+^ CD25^+^ KLRG1^-^ T cells enriched in other Treg subpopulations were analyzed for comparison (Figure 2A).

**Figure 2.**
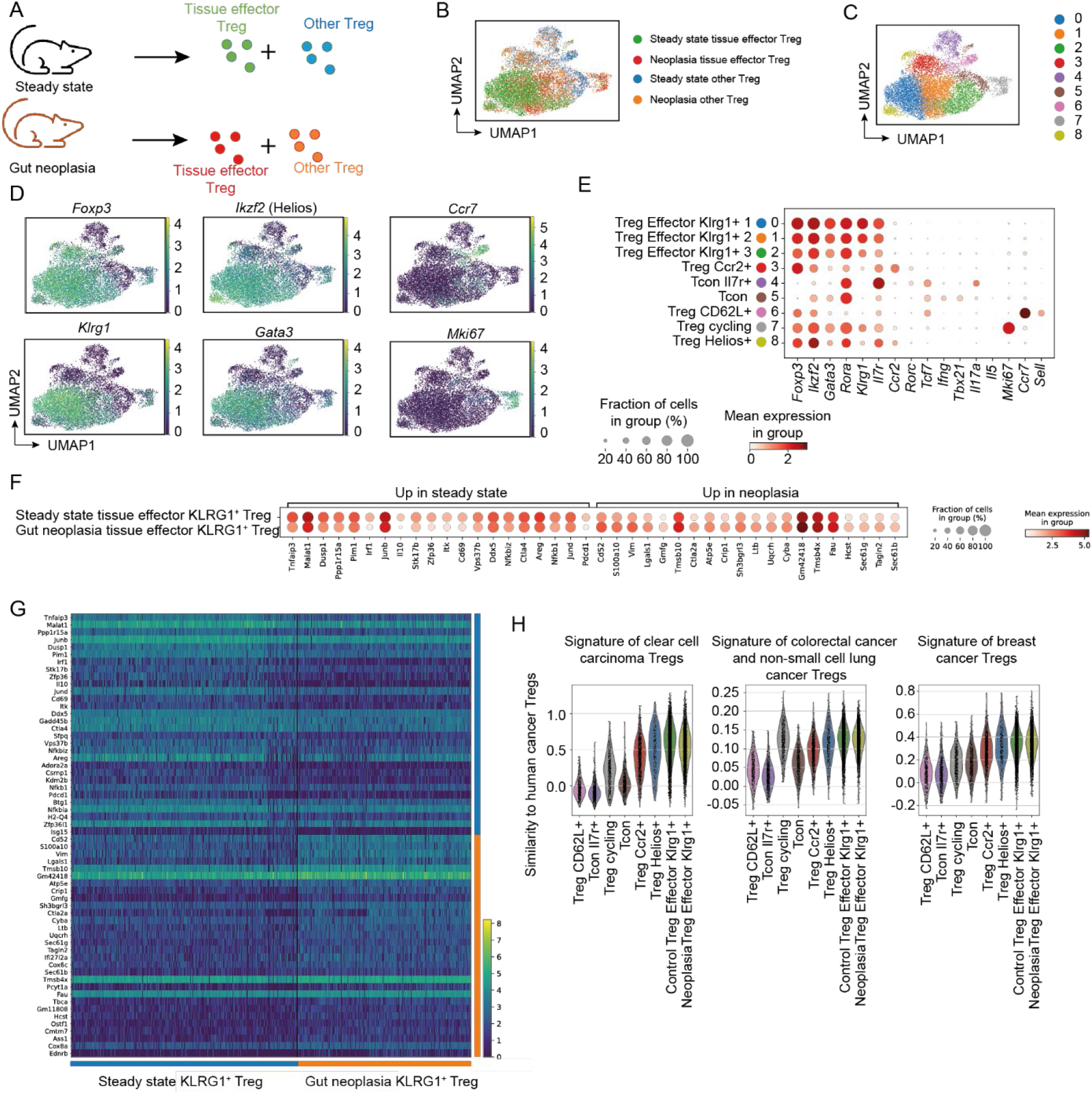
scRNA-seq of neoplasia-responding Tregs. (A) Experimental design. Small intestinal CD25^+^ CD4^+^ T cells from a steady state control and a neoplastic mouse were sorted into “effector tissue Treg” and “other Treg” based on KLRG1 expression and processed for scRAN-seq. (B-D) UMAP of CD4^+^ T cells from the scRNA-seq analysis showing individual samples (B), Leiden cluster analysis (C) and expression of selected genes (D). (E-F) Dot plot (E) showing the signature genes of clusters identified in (B) (F-G) Dot plot (F) and heatmap (G) showing genes in the KLRG1^+^ population (merged clusters 1, 2 and 3) that are differentially expressed between Treg isolated from steady state or neoplastic small intestine. (H) Violin plot showing the similarity score of the identified clusters with published human tumor Treg signatures. Clear cell carcinoma signature from ^14^; CRC/NSCLC (Colorectal cancer/non small cell lung cancer) signature from ^11^; breast cancer signature from ^10^. Control and neoplastic Treg Effector KLRG1^+^ identified as in panels (F-G).

Leiden clustering of processed single cell transcriptomes identified seven *Foxp3*^+^ Treg clusters (clusters 0-3 and 6-8) and two *Foxp3*^*-*^ conventional CD4^+^ T cell (Tcon) clusters (clusters 4 and 5) (Figure 2B-C). In line with the experimental setup, the three most abundant clusters (clusters 0-2) were *Klrg1*^+^ *Gata3*^+^ effector tissue Tregs (Figure 2D-E). *Klrg1* was also expressed by a cluster of proliferating Treg (cluster 7), identified by *Mki67* expression. In addition, we could identify the other main gut Treg clusters: a cluster of lymphoid Treg (cluster 6), characterized by high levels of *Ccr7* and *Sell* (encoding CD62L), a cluster of tissue non effector Tregs expressing *Ikzf2* (encoding Helios) but no *Ccr7* or *Klrg1* (cluster 8), and a cluster characterized by expression of *Ccr2 and Rorc* (encoding RORγt) corresponding to microbiota-dependent Treg (cluster 3) (Figure 2D-E).

Transcriptomic analysis did not identify consistent heterogeneity among the three effector tissue Treg clusters (Figure 2E,S2A, table S1), which a signature including *Klrg1, Gata3, Tnfrsf9*, (encoding CD137) and *Cd83* very prominent in clusters 0 and 1 (Figure 2F, S2, table S1). CD83 has been implicated in tissue Treg homeostasis^17^. Effector tissue Tregs also expressed Amphiregulin (*Areg*) (Figure S2B, table S1), an effector Treg cytokine that promotes tissue repair ^18^. Given the similarities, we merged the three KLRG1^+^ Treg clusters, corresponding to effector tissue Tregs, for further analysis.

To more deeply characterize the effects of gut neoplasia on Tregs, we compared the effector tissue Tregs from neoplastic gut versus steady state gut. This analysis revealed that, despite inducing strong proliferation, intestinal neoplasia did not strongly affect the Treg transcriptome beyond small reductions in some genes such as *Il10* or *Pdcd1* (encoding PD-1) (Figure 2F-G and Table S2). To assess whether the transcriptomes of mouse effector tissue Tregs are coherent with the phenotype of human tumor Tregs, we extracted transcriptomic signatures from published human tumor Treg data ^10,11 14^ (Table S3) and plotted them against our samples (Figure 2H). Effector tissue Treg clusters from both steady state and neoplastic gut scored similarly high when compared to the three independent human datasets, suggesting that the signature currently identified as “human tumor Treg” signature reflects an enrichment in effector tissue Tregs in the tumor environment rather than upregulation of “tumor markers” in Treg populations.

### Epithelial neoplasia induces a strong clonal expansion of private effector tissue Tregs

To determine changes at clonal levels, we analyzed Treg clonality in our scTCR-Seq data. Effector tissue Tregs from both steady state and neoplastic intestine showed oligoclonality as per standard definition (more than 3 cells per clonotype) (Figure 3A). However, clonal size display indicated a much higher clonotype size in effector tissue Treg from neoplastic gut (Figure 3B). Indeed, KLRG1^+^ Tregs were dominated by a smaller number of expanded clones compared to their KLRG1^-^ counterparts, even in steady state (Figure 3C). The difference was even more pronounced after neoplasia, where massively expanded individual effector tissue Treg clones accumulated (Figure 3C). The stark increase in a few clonotypes should lead to a decrease in general clonal diversity. We verified this effect by sequencing the TCRα chain in an independent experiment that confirmed that intestinal neoplasia reduces the diversity of effector tissue Tregs, but not of control KLRG1^-^ Tregs (Figure 3D). Notably, none of the effector tissue Treg expanded clones were appreciably expanded in other mice (Figure 3E), indicating that the clonotypes that expand in response to tumoral changes are private to each mouse. Accordingly, scTCR-Seq data showed no overlap in the expanded clonotypes between control and neoplastic mice (Figure 3F). Most highly-expanded clones contained effector tissue Tregs, dividing cells and a small fraction of non-effector tissue Tregs, although there were also a few clones formed primarily by Tcon cells (Figure 3F).

**Figure 3.**
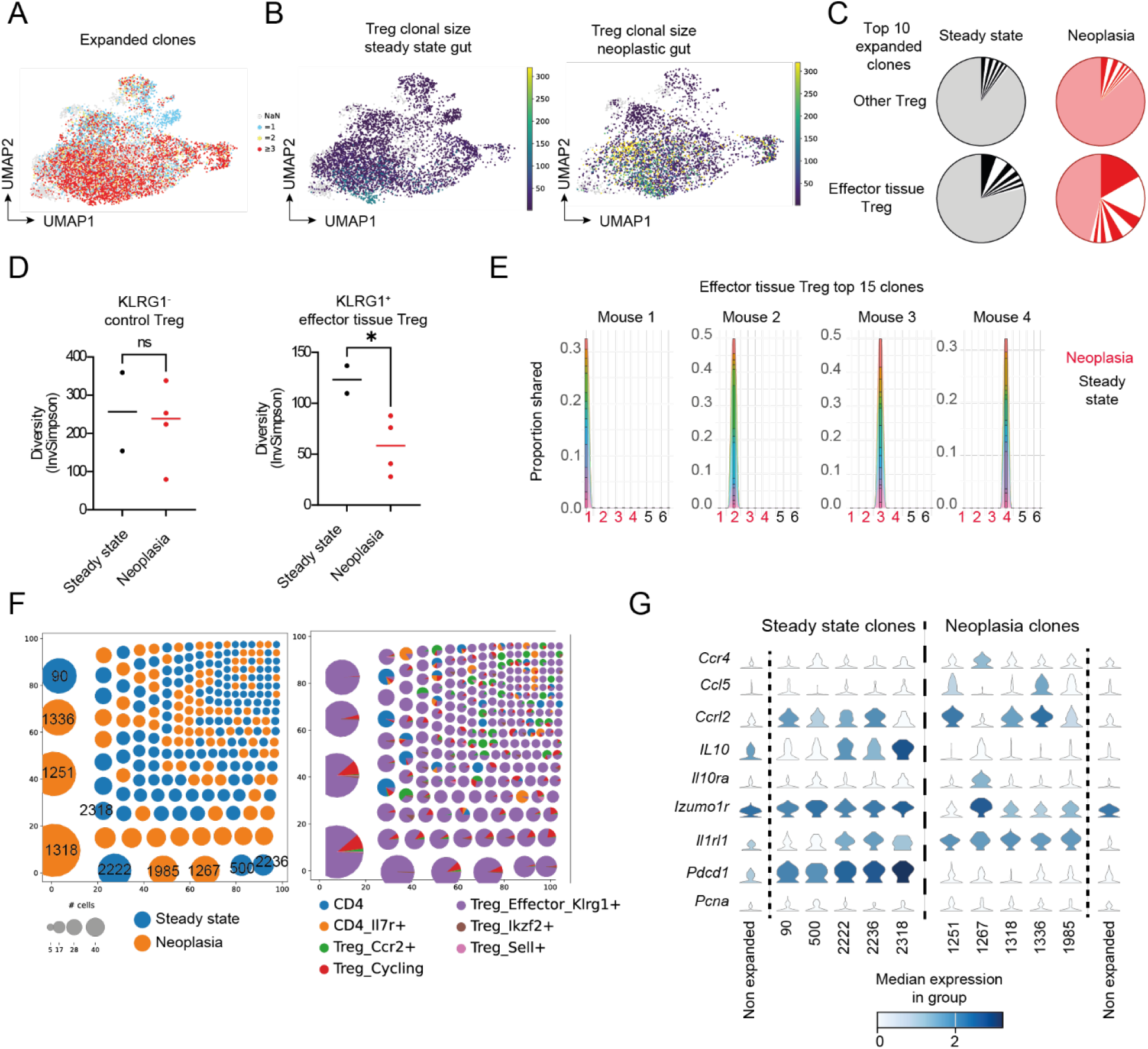
Clonotypic analysis of effector tissue Treg. (A-F and H-K) Analyze data from the combined scRNA-seq and scTCR-seq data depicted in Figure 2 (A-B) UMAPs showing abundance of TCR sequences in Treg as per classical definition (oligoclonality when clonotype size ≥ 3, A) and with a heatmap indicating clonotype size (B). (C) Relative frequencies of the 10 most abundant clones in each sequenced sample. (D-E) Treg from proximal small intestine of individual mice were sorted to either effector tissue (KLRG1+) or other (KLRG1-) Treg, and TCR alpha chains were sequenced to identify clonotypes. (D) Clonal diversity as shown by inverse Simpson index of TCR alpha sequences. Each dot in one plot represents an individual mouse. *, p<0.05; ns = non-significant. Samples compared by unpaired Student’s t test. (E) Proportion of the top 15 most expanded clonotypes of KLRG1^+^ Treg from neoplastic mice among all the sequenced samples. Red numbers: Tregs from neoplastic small intestine; black numbers: Tregs from steady state small intestine; For each tissue, effector tissue Treg are shown to the right, control KLRG1-Treg to the left. (F) Distribution of clonotypes identified from scRNA-Seq across samples (left) and clusters (right). ID numbers shown on left for the most abundant clones. (G) Violin plots showing expression levels of selected genes by the 5 most abundant clonotypes from effector Treg cells isolated from steady-state or neoplastic small intestine. Only cells included in the KLRG1^+^ effector cluster were analyzed. The numbers on the x axis correspond to the clone numbers in (F).

It is known that different Treg subpopulations can respond to different immune cues and use different immunoregulatory mechanisms. To determine if single clones inside of one Treg subpopulation can present such heterogeneity, we characterized clone-associated transcriptional signatures of the effector tissue Treg subpopulation. We included only cells mapping to the KLRG1^+^ effector tissue Treg cluster to avoid contamination biases. As controls for the expanded clones, we used “non-expanded Treg” groups including effector tissue Treg with unique TCR sequences in the sc-TCRseq data. We found that most genes were similarly expressed across clones, including the effector Treg signature genes *Ccr8, Klrg1* and *Gata3* (Figure S3). However, some genes were preferentially expressed by specific clones (Figure 3G). These genes included chemokine/chemokine receptor genes such as *Ccr4*, expressed by cells from one neoplastic clone, and *Ccl5*, expressed by some expanded neoplastic clones. Interestingly, the Treg effector cytokine *Il10* and the alpha chain of the IL-10 receptor (*Il10ra*) showed clone-skewed expression. *Il10* was expressed by all five of the top expanded clones in steady state conditions, but by only 1 out of 5 top expanded clones upon neoplasia (Figure 3G). In contrast, *Il10ra* expression showed clonal variability but not a strong condition bias. *Izumo1r*, recently described to be involved in Treg function ^19^, also showed biased clonotype expression (Figure 3G). In contrast the inhibitory receptor PD-1, encoded by *Pdcd1*, showed consistent increased expression in steady state as compared to neoplasia, while *Cd5* (reflecting T cell activation) and *Pcna* (reflecting cell proliferation) also showed similar levels across the clones, irrespective of neoplasia. These data suggest clonotypic differences in effector tissue Treg effector capacities and response to immune cues.

All in all, TCR analysis shows that effector tissue Tregs are oligoclonal in the steady state and respond to neoplasia by a very strong expansion of private clones.

### Effector tissue Treg response to neoplasia is TCR-dependent but independent of IL-33R, CCR8 and CD137

Since neoplasia induces Treg accumulation through oligoclonal expansion, we checked whether TCR signals are required for this process. To this aim, we generated bone marrow chimeras by reconstituting irradiated *Catnb*^*+/lox(ex3)*^ Vil Cre-ERT mice or Vil Cre-ERT-negative mice with CD4^Cre ERT/+^ TCRa^fl/fl^ bone marrow (Figure 4A). In these mice, tamoxifen injection leads to simultaneous induction of gut epithelial cell neoplasia and TCR ablation in a fraction of CD4^+^ T cells. The ablation affects only a fraction of T cells, allowing for direct comparison between TCR+ and TCR KO cells in the same host.

**Figure 4.**
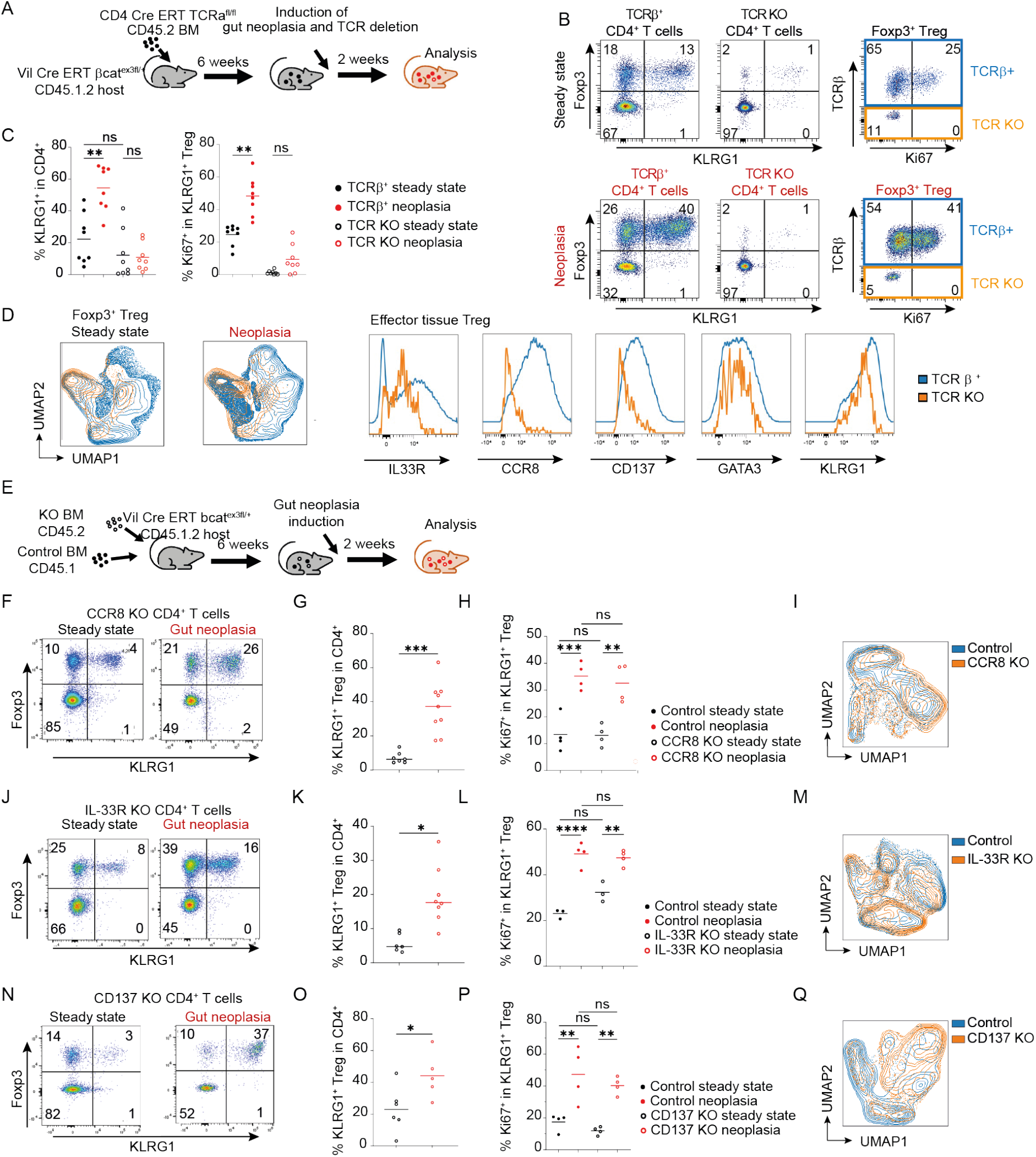
Roles of TCR, IL-33R, CCR8 and CD137 in effector tissue Treg response to neoplasia. (A) Set up of TCRa^fl/fl^ bone marrow chimeras (B-D) Representative FACS stainings (B) and frequencies (C) of effector tissue Treg and proliferating effector tissue Treg among TCR^+^ and TCR^-^ CD4^+^ lymphocytes in steady-state and neoplastic proximal small intestine. (D) UMAP and histogram of FACS analysis of TCR^+^ (blue) and TCR KO (orange) CD4^+^ Treg in steady-state and neoplastic small intestines. UMAP features IL-33R, Granzyme A, KLRG1, Foxp3, CCR8 and Helios on donor Treg (see also Supplemental FACS data). Data show four concatenated mice per group and are representative of at least two independent experiments. (E) Set up of mixed bone marrow chimeras (F-Q) Data show analysis of proximal small intestine. (F-I) Analysis of CCR8 KO /CD45.1 mixed bone marrow chimeras. Representative FACS plots (F) and frequencies (G) of KLRG1^+^ effector tissue Treg among wild-type and CCR8-deficient CD4^+^ T cells from control and neoplastic intestines. (H) Frequencies of dividing KI67^+^ cells among control CD45.1^+^ (solid symbols) and CD45.2^+^ CCR8 KO (empty symbols) CD4^+^ Foxp3^+^ KLRG1^+^ Treg from steady-state (black) and neoplastic (red) gut. G-H data are pooled from at least two independent experiments. (I) UMAP of CCR8-deficient and control CD45.1 KLRG1^+^ Treg. UMAP was performed on 10,000 concatenated donor Treg per mouse isolated from 2 steady state and 2 neoplastic mice. UMAP features IL-33R, Granzyme A, KLRG1, Foxp3, Helios and CD137. Control CD45.1^+^ in blue, CCR8 KO in orange. (J-M) Analysis of IL-33R KO /CD45.1 mixed bone marrow chimeras. Representative FACS plots (J) and frequencies (K) of KLRG1^+^ effector tissue Treg among wild-type and IL-33R-deficient CD4^+^ T cells from control (black) and neoplastic (red) intestines. (L) Frequencies of dividing KI67^+^ cells among control CD45.1^+^ (solid symbols) and CD45.2^+^ IL-33R KO (empty symbols) CD4^+^ Foxp3^+^ KLRG1^+^ Treg from steady-state (black) and neoplastic (red) gut. (M) UMAP of IL-33R-deficient and control CD45.1 KLRG1^+^ Treg. UMAP was performed on 10,000 concatenated donor Treg per mouse isolated from 2 steady state and 2 neoplastic mice. UMAP features Granzyme A, KLRG1, Foxp3, CCR8, Helios and CD137. Control CD45.1^+^ in blue, IL-33R KO in orange. K-L Data are pooled from at least two independent experiments. (N-Q) Analysis of CD137 KO /CD45.1 mixed bone marrow chimeras. Representative FACS plots (N) and frequencies (O) of KLRG1^+^ effector tissue Treg among wild-type and CD137-deficient CD4^+^ T cells from control and neoplastic intestines. (P) Frequencies of dividing KI67^+^ cells among control CD45.1^+^ (solid symbols) and CD45.2^+^ CD137 KO (empty symbols) CD4^+^ Foxp3^+^ KLRG1^+^ Treg from steady-state (black) and neoplastic (red) gut. (Q) UMAP of CD137-deficient and control CD45.1 KLRG1^+^ Treg. UMAP features IL-33R, Granzyme A, KLRG1, Foxp3, CCR8 and Helios on 10,000 concatenated donor Treg per mouse isolated from 2 steady state and 2 neoplastic mice. Control CD45.1^+^ in blue, CD137 KO in orange. N-P Data are pooled from at least two independent experiments. (C, H, L, P): ANOVA followed by Šídák’s multiple comparisons test. (G, K, O): unpaired Student’s t test. *, p<0.05; **, p<0.01; ***, p<0.001; ns, non significant.

We then examined the frequencies of TCR KO cells among different Treg populations. In the steady state intestine, Tregs had a more stringent requirement for TCR than conventional T cells, as assessed by a lower frequency of Foxp3^+^ cells among TCRβ-negative cells (Figure 4B, S4A). TCR ablation completely abrogated Treg cell division under steady-state conditions, as assessed by Ki67 expression (Figure 4B). When we compared the response of TCR-positive and TCR-deficient cells, we found that TCR-deficient Treg failed to divide and did not expand in response to neoplasia (Figure 4B, C). We then characterized the Treg phenotype by multiparametric flow cytometry. UMAP display showed that the remaining TCR-deficient effector tissue Tregs in neoplastic gut had a distinct phenotypic profile (Figure 4D, supplemental FACS data), including lower levels of CCR8 and CD137 than wild-type cells, suggesting that both features are driven by cognate antigen signals. Expression of IL-33R was detectable, although at lower levels than in wild-type cells. KLRG1 expression was only slightly reduced and GATA3 unchanged in TCR-deficient cells (Figure 4D, Supplemental FACS data). Hence, TCR signals are essential for tissue Treg proliferation and expansion to neoplastic cues.

CCR8, IL-33R and CD137 are all part of the effector tissue Treg signature and have been implicated in T cell expansion. CCR8 was the first marker associated with tumor Tregs, and it can promote T cell responses ^20^. IL-33R specifically induces Treg accumulation in the gut ^21^, and CD137 is a costimulatory molecule modulating T cell responses ^22^. We therefore set out to analyze the effect of these molecules on the neoplasia-induced expansion of effector Tregs. To this aim, we reconstituted irradiated *Catnb*^*+/lox(ex3)*^ Vil Cre-ERT host mice co-expressing the CD45.1 and CD45.2 congenic markers with a mixture of CD45.1^+^ wild-type bone marrow and CD45.2^+^ bone marrow from mice deficient for the gene of interest (Figure 4E).

We first analyzed the role of CCR8. We found that CCR8-deficiency did not prevent effector tissue Treg expansion in response to gut neoplasia (Figure 4F-G, S4B), and lack of CCR8 did not affect effector Treg proliferation (Figure 4H). Further, and in line with published data ^14,23,24^, CCR8 deficiency did not affect effector Treg phenotype in neoplastic guts (Figure 4I, supplemental FACS data). Hence, CCR8 does not play an essential role in effector Treg expansion in response to gut neoplasia.

We next turned to IL-33R. IL-33 is upregulated by neoplasia and it promotes effector Treg accumulation in the gut under steady-state conditions ^21^. Accordingly, IL-33R-deficient bone marrow was less efficient in reconstituting the KLRG1^+^ effector tissue Treg compartment as compared to the control Treg compartment (Figure S4B). However, IL-33R-deficient Treg still expanded in response to neoplasia (Figure 4J-M). While the frequency of effector tissue Tregs was reduced among gut steady-state IL-33R KO Treg compared to wild-type Tregs, the fold-increase induced by neoplasia was comparable for IL-33R-deficient and control cells (Figure S4C-D). This indicates that effector tissue Treg accumulation in the gut is dependent on IL-33R under steady-state conditions, but IL-33R does not mediate neoplasia-induced effector Treg accumulation. FACS analysis showed that the phenotype of neoplasia effector tissue Tregs was largely independent on IL-33R (Figure 4M), with differences in the UMAP profile being due to slightly lower GATA3 and KLRG1 expression in IL-33R-deficient effector Treg (Supplemental FACS data). Hence, despite playing an important role in effector tissue Treg accumulation in the steady state, IL-33R does not mediate neoplasia-driven Treg accumulation and is largely dispensable for the effector tissue Treg signature.

We then assessed the role of the costimulatory protein CD137, a potential candidate for immune checkpoint treatment that, like PD-1 and CTLA-4, is also highly expressed by effector Tregs. In contrast to IL-33R-deficient cells, CD137 KO Treg expanded more efficiently than control Treg under steady-state conditions (Figure S4B). CD137 deficiency did not impair effector Treg expansion or division in response to intestinal neoplasia (Figure 4N-P) nor strongly affected the phenotype of KLRG1^+^ effector Treg (Figure 4Q). Minor differences between control and CD137 KO effector Tregs in the UMAP profile were accounted for by higher expression of IL-33R and KLRG1 in the CD137 KO effector Treg population (Supplemental FACS data). This observation, together with the increased accumulation of CD137-deficient effector tissue Tregs, suggests that CD137 may exert cell-intrinsic inhibitory effects on Tregs. Nevertheless, CD137 was not required for neoplasia-induced effector Treg response or for the effector tissue Treg signature.

CD83 is required for KLRG1^+^ Tregs^17^ and is part of the effector tissue Treg signature. We checked its role on neoplasia-driven Treg accumulation using Foxp3^YFP-Cre^ CD83^fl/fl^ bone marrow to generate chimeras as in Figure 4E. The widely used Foxp3^YFP-Cre^ allele contains an IRES ^YFP-Cre^ fusion protein inserted after the *Foxp3* locus, which induces Treg-specific CD83 deletion and allows Cre expression to be tracked by the detection of YFP ^25^. Measuring YFP by flow cytometry in these cells was not compatible with the intracellular staining protocol required for Foxp3 detection. Therefore, to check for deletion efficiency, we compared frequencies of YFP^+^ with frequencies of Foxp3^+^ cells in parallel stainings (Figure S4E-F). We found that almost all Foxp3^YFP-Cre^ CD83^fl/fl^ effector tissue Tregs were deletional escapees, suggesting a profound role for CD83 in effector tissue Treg homeostasis. In contrast, neoplasia-induced accumulation of effector tissue Tregs accumulated was independent of CD83 (Figure S4G).

All in all, our data show that effector tissue Treg accumulation in response to neoplasia is antigen driven, but independent of mechanisms that affect tissue Treg accumulation in the steady-state such as IL-33 and CD83.

### Treg expansion in response to intestinal neoplasia relies on cell-intrinsic GATA3

Since most KLRG1^+^ Tregs express the transcription factor GATA3, we then analyzed its requirement for effector tissue Treg homeostasis. Lack of GATA3 causes embryonic lethality, so we used the published GATA3^fl/fl^ Foxp3^YFP-Cre^ mouse strain (from here on referred to as GATA3 KO) to specifically deplete GATA3 in Treg ^26,27^. It has been previously shown that GATA3 KO Treg have lower levels of Foxp3 and reduced KLRG1 expression, but the effect of GATA3 on different Treg subsets has not been addressed ^26-28^.

We found that GATA3 KO mice had normal levels of KLRG1^+^ Treg in spleen compared to mice with GATA3-heterozygous Tregs (Figure 5A and S5A). In contrast, despite overall higher Treg frequencies (^26^ and Figure S5B), absence of Treg-intrinsic GATA3 expression led to a striking reduction in effector tissue Tregs in the gut (Figure 5A). This reduction was not simply due to downregulation of KLRG1, since the effector tissue Treg population was also absent when IL-33R or CD137/CCR8 were used as markers for effector tissue Tregs (Figure 5B). Of note, IL-33R expression on KLRG1 negative Tregs was not reduced, indicating that GATA3 expression is dispensable for IL-33R expression on Tregs, but necessary for the intestinal effector tissue Treg subset. Effector tissue Tregs account for a sizeable proportion of all intestinal Tregs. Therefore, if GATA3 were required for survival of effector tissue Treg, we would expect lower frequencies of Helios^+^ Treg in the gut. However, the frequency of Helios-expressing Tregs was not decreased in the absence of GATA3 (Figure 5C). This result indicates that GATA3 deficiency does not lead to the death of Helios^+^ GATA3^+^ Treg, but rather to a lack of differentiation into effector tissue Tregs.

**Figure 5.**
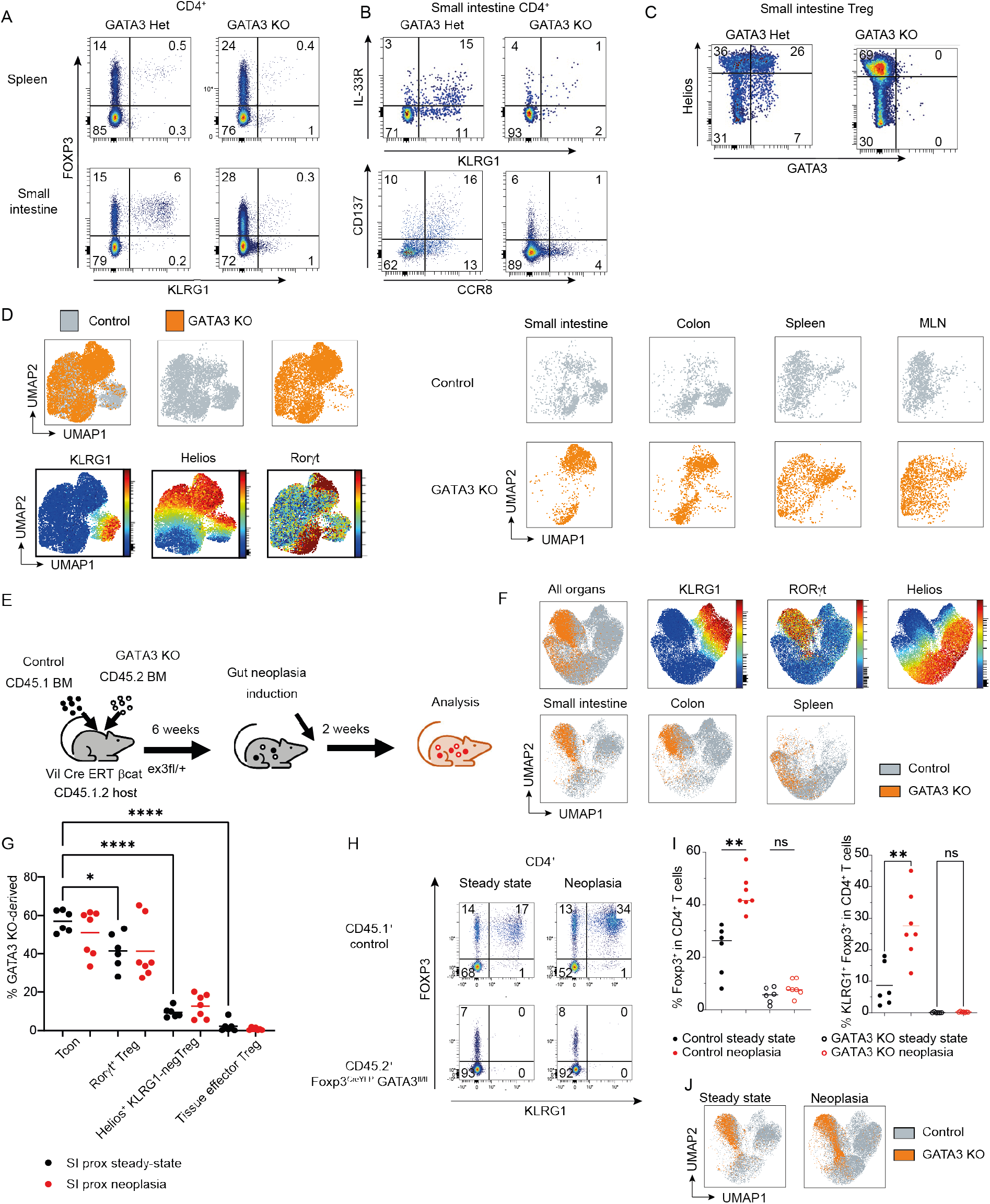
Role of GATA3 in in effector tissue Treg response to neoplasia. (A-C) Representative FACS plots of CD4^+^ T cells (A) and CD4^+^ Foxp3^+^ Treg (B, C) from the indicated organs from GATA3^fl/+^ Foxp3^YFP-Cre^ BL/6 and GATA3^fl/fl^ Foxp3^YFP-Cre^ mice. (D) UMAP of Treg from control CD45.1 (grey) and GATA3^fl/fl^ Foxp3^YFP-Cre^ (orange) mice from the indicated organs. UMAP was performed on IL-33R, GzmA, CTLA-4, PD-1, KLRG1, RORγt, Helios, CD137, CD44 and CD25; see also Figure S5B. (E) Set up of GATA3 KO mixed bone marrow chimeras. (F-J) Analysis of the mice in (E) (F) UMAP of control CD45.1^+^ (grey) and GATA3^fl/fl^ Foxp3^YFP-Cre^ CD45.2^+^ (orange) Treg from the indicated organs of mixed bone marrow chimeras. UMAP was performed on IL-33R, CTLA-4, PD-1, KLRG1, CCR8, RORγt, Helios and CD44; see also Figure S5C. (G) Frequencies of GATA3^fl/fl^ Foxp3^YFP-Cre^ –derived cells among conventional CD4^+^ Foxp3neg, Rorgt^+^ Treg, Helios^+^ KLRG1neg Treg and Helios^+^ KLRG1^+^ effector Tregs. (H-I) Representative FACS plots (H) and frequencies (I) of KLRG1^+^ effector tissue Treg among wild-type and GATA3^fl/fl^ Foxp3^YFP-Cre^ CD4^+^ T cells from control and neoplastic intestines. (J) UMAP as in (F) comparing Treg from neoplastic small intestine to the ones from steady state small intestine from (F).

UMAP display of flow cytometry analysis indeed showed that the intestinal effector tissue Treg population is virtually missing in the absence of GATA3 expression by Tregs as compared to control mice, whereas we detected a Helios^+^ RORγt^+^ double positive population in the small intestine and colon of GATA3 KO that is not conspicuous in normal mice ^29,30^ (Figure 5D). To determine if the effect of GATA3 on effector tissue Tregs was cell intrinsic, and to check the response of GATA3-deficient Tregs to neoplasia, we generated mixed bone marrow chimeras using CD45.1 and GATA3 KO bone marrow (Figure 5E). We found a strong effect of GATA3 on all Treg subtypes except Helios^-^ RORγt^+^ Tregs. In competition under steady state conditions, GATA3 KO Tregs did not strongly contribute to the effector tissue Treg pool in lymphoid organs nor intestinal tissue (Figure 5F and S5C). Importantly, GATA3 KO Tregs were also strongly reduced in the Helios^+^ KLRG1^-^ non-effector tissue Treg pool. This observation suggests that, while GATA3 is essential for effector tissue Tregs, it also contributes to the competitive fitness of Helios^+^ Tregs (Figure 5F). In contrast, the Helios negative RORγt^+^ Treg population was mostly independent of GATA3. We confirmed these observations by checking the contribution of GATA3 KO cells to donor-derived Treg subsets in the small intestine. Indeed, cells derived from GATA3 KO bone marrow were readily present in the Helios^-^ RORγt^+^ Treg compartment (Figure 5G); the contribution of GATA3^-^deficient cells to this population was just slightly lower than for conventional Tcon. In contrast, GATA3-deficient Tregs were strongly reduced among Helios^+^ non effector Tregs, and they were virtually absent from the KLRG1^+^ Treg pool (Figure 5G). Intestinal neoplasia did not affect the frequencies of GATA3-deficient Tregs in the different subsets (Figure 5G). The rare KLRG1^+^ Tregs coming from the GATA3 KO bone marrow were, in fact, GATA3-expressing deletion escapees (Figure S5D).

In contrast to other tested KO, GATA3 KO Tregs failed to accumulate in response to neoplasia (Figure 5H-I). Multiparametric flow cytometry showed that the phenotype of GATA3-deficient Tregs was not substantially altered under intestinal neoplasia (Figure 5J), except for a small but significant increase in Helios^+^ RORγt^+^ cells in response to neoplasia (Figure S5E). This increase might indicate that KO Tregs respond to neoplasia, but that in the absence of GATA3 they adopt a Helios^+^ RORγt^+^ phenotype that cannot sustain Treg accumulation in competitive settings. We were unable to address the expansion of GATA3 KO Tregs in non-competitive settings because of the strong accumulation of GATA3^+^ deletional “escapees” in this scenario (Figure S5F). Hence, our data suggest that GATA3 expression is not only required for gut effector tissue Tregs, but is also essential for Treg accumulation in response to intestinal neoplasia.

### Tumor-induced Treg expansion and Treg pro-tumoral function rely on biallelic GATA3 expression

Once we established a role for GATA3 in effector Tregs, we assessed if its function is sensitive to gene dosage, as shown for other GATA3 functions ^31,32^. Indeed, mice with heterozygous GATA3 expression in Tregs (GATA3 het) showed a significant reduction in gut effector tissue Tregs, although not to the extent of GATA3 KO, and normal levels of Foxp3 (Figure 6A-B and S6A). In addition, we observed a specific reduction in effector tissue Tregs driven by Foxp3^YFP-Cre^ expression even in the absence of GATA3 modifications, which was not as profound as in GATA3 het mice (Figure 6C-D).

**Figure 6.**
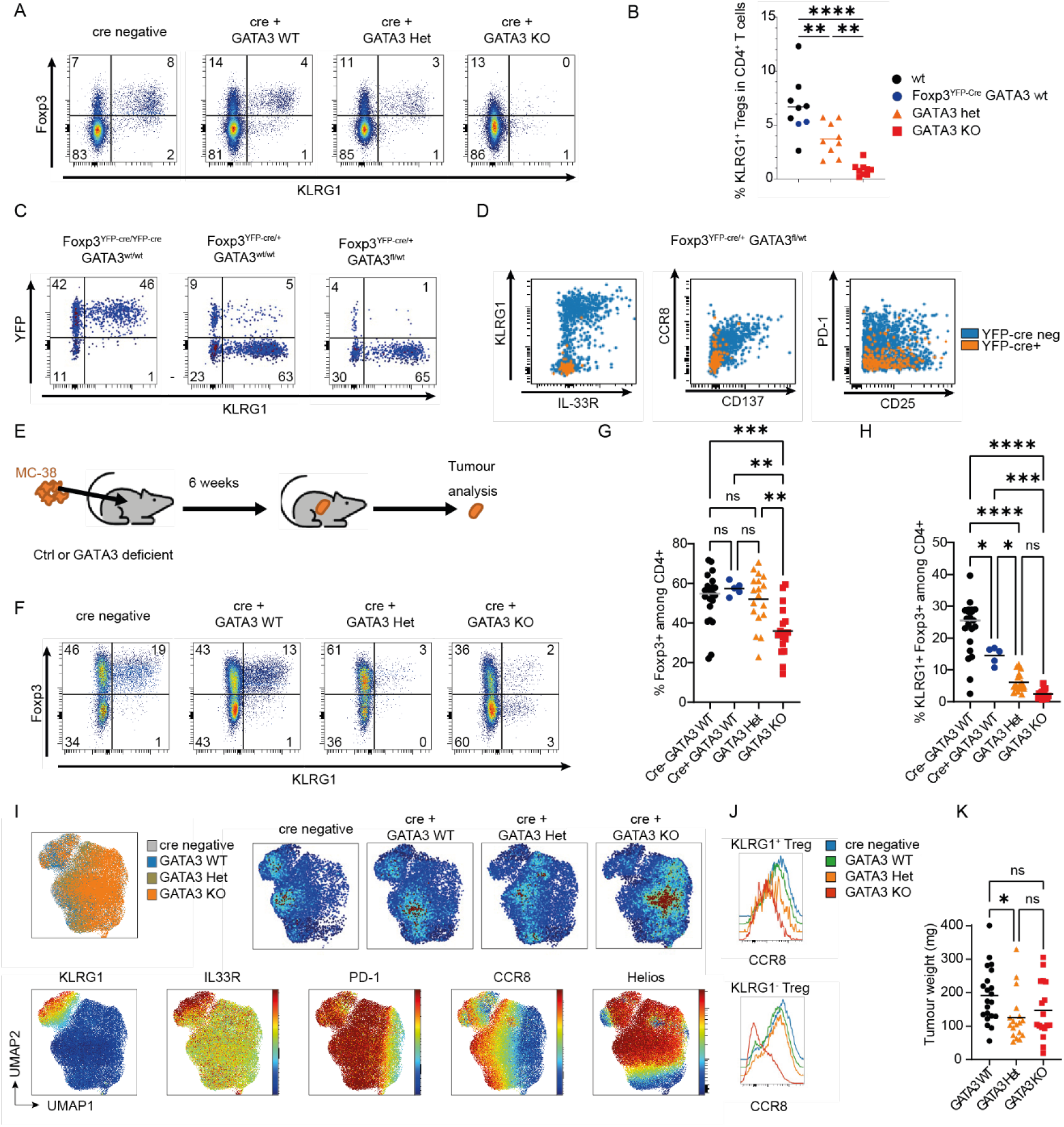
Mono-and biallelic effects of GATA3 in Treg response to tumors. (A-B) Representative FACS plots (A) and frequencies (B) of tissue Treg in proximal small intestine from control mice or Foxp3^y/YFP-Cre^ mice with one, two or no functional GATA3 alleles in Treg. The FACS plots are gated on CD4^+^ T cells. (C) Representative FACS plots of CD4^+^ CD25^+^ T cells from colon of a female expressing Foxp3-^YFP-Cre^ on both (Foxp3^YFP-Cre/YFP-Cre^ GATA3^+/+^) or only on one X chromosome (Foxp3^+/YFP-Cre^) and biallelic or monoallelic GATA3 on YFP^+^ cells. (D) Representative FACS plots showing overlayed Cre YFP negative (blue) and Cre YFP^+^ GATA3 Het (orange) CD4^+^ CD25^+^ T cells from proximal small intestine from a Foxp3^+ /YFP-Cre^ GATA3^fl/+^ female. (E-K) Analysis of cells infiltrating ectopically transplanted tumors. MC38 cells were subcutaneously injected into wild-type mice, wild-type mice expressing Foxp3^YFP-Cre/YFP-Cre/y^ or into mice with wild-type (GATA3^+/+^), heterozygous (GATA3 ^+/fl^) or KO GATA3 in Foxp3^YFP-Cre/YFP-Cre^ Treg. Tumors were analyzed 3 weeks after transplant. (E) Experimental setup (F-H) Representative FACS plots (F) and frequencies of total (G) and tissue-effector Treg (H) among CD4^+^ T cells in tumors. FACS plots are gated on CD4^+^ T cells. (I) UMAP of Foxp3^+^ Treg in the tumors of the mice in (H-I). UMAP was performed on IL-33R, GzmA, Sca1, CTLA-4, PD-1, KLRG1, CCR8, RORγt, Helios, CD137, GATA3, CD25. (J) CCR8 expression on tissue Treg, gated as KLRG1^+^ Foxp3^+^, and other Treg, gated as KLRG1^-^ Foxp3^+^, from MC38 tumors. (K) Weights of the transplanted tumors Data are pooled from 5 different experiments. Statistics: ANOVA followed by Šídák’s multiple comparisons test. *, p<0.05; **, p<0.01; ***, p<0.001; ****, p<0.0001; ns, non significant.

To account for the multiple variables affecting effector tissue Tregs in the GATA3 KO model, we used mice with four genotypes to check if GATA3 is also relevant for Treg accumulation in established tumors. For that, we turned to a widely used ectopic model of intestinal tumor transplant (Figure 6E). In this system, subcutaneous transplant of the mouse colonic cell line MC38 leads to an accumulation of Tregs, especially KLRG1^+^ effector tissue Tregs ^33^. We compared four genotypes: mice expressing ^YFP-Cre^ (both *Foxp3*^*YFP-cre/y*^ males or *Foxp3*^*YFP-cre/YFP-cre*^ females) and either one, two or no floxed GATA3 alleles (GATA3 het, GATA3 KO, Cre+ GATA3 wt), and mice without *Foxp3*^*YFP-Cre*^ (Cre-). We found that the frequency of total Tregs was reduced in tumors from mice with Treg-specific biallelic GATA3 deletion, but not in mice with monoallelic GATA3 expression or in *Foxp3*^*YFP-Cre*^ *Gata3*^*+/+*^ mice as compared with control mice (Figure 6F-G). In contrast, mice with monoallelic GATA3 expression showed a specific reduction in tumor infiltration of KLRG1^+^ effector tissue Tregs, which was comparable to that in GATA3 KO mice (Figure 6F, H). While *Foxp3*^*YFP-Cre*^ + GATA3 wt mice also showed a reduction of effector tissue Tregs in the tumor compared to *Foxp3*^*YFP-Cre*^ - mice, it was not as profound as the one observed in GATA3 het mice. In agreement with a cell-intrinsic role for GATA3 in tumor Treg accumulation, analysis of tumors transplanted into Foxp3^+/YFP-Cre^ GATA3^+/fl^ mice showed that most tumor Treg were GATA3 WT (Figure S6B). Hence, biallelic GATA3 expression is required for the accumulation of effector tissue Tregs in established tumors. Tumor-infiltrating GATA3 KO Tregs showed a distinct surface phenotype from their wild-type counterparts, with lower CCR8 expression in all Tregs (Figure 6 I-J, Supplemental FACS data). In contrast, in agreement with phenotypic data under steady state, the phenotype of GATA3 Het Tregs was similar to the Tregs from GATA3-sufficient mice (Figure 6I, Supplemental FACS data).

We wanted to assess if effector tissue Tregs have specific pro-tumoral functions, or if it is the frequency of total Tregs that counts, irrespective of the Treg phenotype. We therefore assessed tumor weight in the different mutant mice (Figure 6K). Indeed, the tumors in GATA3 Het mice were smaller compared to the ones harboring wild-type Tregs, suggesting that effector tissue Tregs have non-redundant functions in the pro-tumoral immune niche.

All in all, our results demonstrate that GATA3 plays a dual role in Tregs. Monoallelic GATA3 expression in Tregs can rescue Foxp3 expression and Treg accumulation, but it cannot restore effector tissue Treg frequencies. Accordingly, tumor growth was impaired despite unchanged frequencies in total Tregs. Our data hence reveal a non-redundant role for GATA3-driven effector tissue Tregs in favoring tumor growth.

## Discussion

Tregs are key cancer promoters ^14^, but it is unclear how the characteristic tumor Treg compartment arises. Here we show evidence that Tregs respond to early tumoral changes by a rapid GATA3-dependent clonal accumulation of effector tissue Tregs. We used a mouse model of gut neoplasia to characterize Treg accumulation, but the phenotype and oligoclonality of Tregs in our model of intestinal neoplasia resemble the ones found in models of transplanted tumors and in human patients. The findings suggest that the Treg population that gets established during the first neoplastic events persist during cancer development. Our results show that Tregs can respond to early events in intestinal tumor development by GATA3-dependent clonal expansion of tissue-effector Tregs, without substantially altering their Treg phenotype.

Our observations in this model confirm that an early molecular event alone is enough to drive an oligoclonal accumulation of effector tissue Tregs, similar to the Treg accumulation observed in human patients and different preclinical models ^9,34^. While a phenotype associated with tumor Tregs has long been proposed based on the common features of Tregs in different organs ^10,11,13^, these comparisons were often population-based or used lymphoid Tregs as controls. The use of a neoplastic model allows a direct comparison with tissue Tregs from healthy gut. We show that activation of the epithelial Wnt pathway, an early event in the oncogenic progression of intestinal tumors ^8^, induces the local accumulation of effector tissue Tregs. This accumulation was driven by TCR signals and was independent of IL-33R, a major driver of effector tissue Treg accumulation during steady state that induces effector tissue Treg proliferation *in vitro* ^9,21^. Indeed, previous reports suggest that IL-33R promotes the accumulation of Tregs in tumors ^12,35-37^. Our results indicate that IL-33R plays a role in effector tissue Treg accumulation before tumorigenesis, which fits with its expression in precursors of effector tissue Tregs in lymphoid organs ^5^, but is dispensable for the actual neoplasia-induced accumulation of Tregs. We observed a similar dichotomy with CD83, which plays an essential role in effector tissue Treg accumulation during steady state ^17^ but is dispensable for effector tissue Tregs in response to neoplasia.

In contrast, our results suggest an important role for TCR signals in cell division and effector tissue Treg phenotype. TCR controls CD137 and CCR8 expression, whereas IL33R and KLRG1 expression on effector tissue Tregs are enhanced by GATA3, which also enhances CCR8. Our scRNA-seq analysis also suggests that the effector Treg phenotype is enhanced during activation, leading to a stronger phenotype in acutely activated Tregs in preclinical models as compared to chronic cancer models.

Our scRNAseq analysis shows that “tumor Treg” signatures most likely reflect a transcriptomic signature shared by effector tissue Tregs from both tumors and healthy tissue, with the fraction of effector tissue Tregs swiftly increasing during tumorigenesis. The specific phenotype of tumor Tregs is an attractive target for anti-tumoral therapy, and current research is identifying exciting Treg-based strategies to fight cancer ^1,38-40^. Knowing that effector tissue Tregs share the same phenotype will help predict side effects of such therapies. it will be important to check how much the effector tissue Treg populations outside of tumors are affected by such treatments. Human tumor Tregs are enriched for the markers we find in mouse effector Tregs ^10,11,13,14^, which suggests similar mechanisms may shape tumor Tregs in both species. However, effector tissue Treg markers were present in human tumors at lower frequencies than in the mouse models we examined. This could be due to several factors. On the one hand, mice and humans present species-specific differences. For example, gut human Tregs are much less frequent as compared to their counterparts in the mouse intestine, even in the steady state ^15^. On the other hand, our model is conceived to induce a strong synchronized response in Tregs. It may be that in human tumors, which typically develop over years, the Treg response includes cells at different activation stages. In agreement with a role for kinetics in the behavior of effector tissue Tregs in tumors, we found that the frequency of dividing Tregs drastically drops four weeks after initiation of neoplasia.

An important element is the striking clonal expansion induced by neoplasia. Tumors are associated with Treg oligoclonality, as are effector tissue Tregs ^5,14^. However, our data show that early tumoral changes are enough to induce a massive expansion of some clones of effector tissue Tregs. The rate of expansion seems to slow down at four weeks, suggesting that the events during initial neoplastic changes are key to shape the immunosuppressive repertoire in tumors. Another possibility, to be addressed by future research, is whether different “clonal waves” occur, where Treg clones are replaced by newly arriving clones. The stability of the effector tissue Treg phenotype inside of a clone also remains to be tested ^41,42^. As for most tumoral responses, the antigens driving effector tissue Tregs in our neoplastic model are unknown. Since this model does not involve the generation of neoantigens, we assume that the response is probably driven by self-antigens.

Analysis of mice with GATA3-deficient Tregs previously failed to identify the absence of the effector tissue Treg subset ^26-28,43^. This is probably due to the use of mice with heterozygous GATA3 Tregs as controls, which suffer from the combined effects of GATA3 haploinsufficiency and of the specific adverse effects of *Foxp3*^*YFP-Cre*^ expression on effector tissue Tregs. Such adverse effects are probably not unique to the *Foxp3*^*YFP-Cre*^ allele ^25^. Indeed, an independently generated *Foxp3*^*GFP-Cre*^ line ^44^ also shows negative effects on Tregs, underlining the need to analyze different genotypes as controls. Considering the four different genetic variants of GATA3 and Foxp3^YFP-Cre^ used in this study, we conclude that GATA3 is essential for effector tissue Tregs, and that monoallelic GATA3 is not enough to restore the fitness of the intestinal effector tissue Treg compartment.

The model of tamoxifen-induced neoplasia presents the advantage that the neoplastic changes are largely independent of the immune response, which allows dissecting the effects of neoplasia on Tregs. In contrast, this model does not allow the assessment of the effects of Tregs on tumor development. We therefore used a well-described model of heterotopically transplanted tumors with Treg accumulation to identify functional effects of effector tissue Tregs ^33^. Analysis of cell-intrinsic effects allowed us to dissect the effects of GATA3 KO and monoallelic GATA3. As previously shown ^26-28^, GATA3 KO affected all Treg, through mechanisms including a decreased level of Foxp3 expression per cell. GATA3-KO Tregs showed reduced accumulation in tumors, while monoallelic GATA3 restored total Treg frequencies. Strikingly, monoallelic GATA3 expression was not sufficient to restore effector tissue Treg accumulation in MC38 tumors. The reduction in effector tissue Tregs was a compound effect of Foxp3 ^YFP-Cre^ expression and lack of biallelic GATA3. Importantly, reduction in effector tissue Treg frequencies led to smaller tumor sizes despite similar total Treg levels. This indicates that effector tissue Tregs play a role in tumor growth that cannot be fulfilled by other Treg subsets.

In summary, our data show that GATA3 is essential for a Treg population non-redundantly promoting tumor growth, and further supports the notion that early tumorigenic events set the stage for oligoclonal Tregs to establish an immunosuppressive environment.

## Supporting information

Supplemental FACS data

## Supplemental information

Document S1. Figures S1-S6

Table S1

Table S2

Table S3

Supplemental FACS data

## Acknowledgements

We thank Vuk Cerovic for helpful discussion and editorial contributions. The work was supported by the Flow Cytometry Facility, the Transgenic Facility and by the Genomics Facility, core units of the Interdisciplinary Center for Clinical Research (IZKF) Aachen within the Faculty of Medicine at RWTH Aachen University. This work was funded by the German research foundation (DFG) Project-ID 449790317 to A.I., Project-ID 403224013 – SFB 1382 (B08 to A.I. and U.N., B06 to O.P) and by the START-Program of the Faculty of Medicine RWTH Aachen University.

We thank F. Powrie for generously providing the IL33R KO mice, Y. Belkaid, N. Bouladoux and J. Zhou for kindly providing the Foxp3^y/YFP-cre^ GATA3^fl/fl^ mice and S. Wirtz for providing CCR8 KO mice. We thank Trong-Hieu Nguyen for help with data submission, and J. Steitz for help with the MC-38 model. We thank K. Vukovic, R. Grychtol and P. Ergezinger for excellent technical support.

## Declaration of Interests

The authors declare no competing interests.

## Material and Methods

### 1. Animal experiments

All mice used in experiments were housed under a 12 h light/dark cycle. Experiments were performed with mice of both sexes that were at least 8 weeks old. The following mouse strains on the C57BL/6 background were bred and kept in the animal facility of the Uniklinik RWTH Aachen under specific pathogen-free conditions:

Wild-type C57BL/6/J (wt), B6.SJL-*Ptprc*^*a*^ *Pepc*^*b*^/BoyJ (named CD45.1), *Catnb*^*+/lox(ex3)*^ B6.Cg-Tg(Vil1-cre/ERT2)23Syr/J(named *Catnb*^*+/lox(ex3)*^ Vil Cre-ERT) ^8,45^, TCRα^fl/fl^ CD4^CreERT2/+^ (TCR KO) ^3^, B6;129-*Il1rl1*^*tm1Anjm*^/J (IL-33R KO) (kindly provided by Fiona Powrie, University of Oxford) ^46^, GATA3^tm1Jfz^ B6.129(Cg)-Foxp3^tm4(YFP/icre)Ayr^/J (GATA3-fl Foxp3^YFP-Cre^) (generously provided by Yasmin Belkaid, NIAID Bethesda) ^27^, Ccr8^tm1Lira^ (named CCR8 KO) ^20^ (kindly provided by Stefan Wirtz, Uniklinikum Erlangen) and B6;129S2-Tcra^tm1Mom^/J (TCRa KO) were kept and bred at the animal facilities of the RWTH University Clinic of Aachen. *Tnfrsf9*^*tm1Byk* 22^ (named CD137 KO mice) were kept and bred at the animal facilities of the University Clinic of Hannover. Cd83^tm1.1Lnjt^ (CD83 KO) mice ^17^ were kept and bred at the University of Erlangen. *Cdh1*^*fl/Ncad*^ Vil Cre^+^ cadherin-switched mice ^47^ were kept and bred at the animal facilities of the Max Planck Institute for Immunobiology and Epigenetics in Freiburg.

For use in experiments, VilCreERT2^+/-^ β-cat^ΔEx3/ ΔEx3^ males were bred with CD45.1^+^ females to give rise to VilCreERT2^+/-^ β-cat^ΔEx3/+^ CD45.1^+^ CD45.2^+^ animals.

Animals were sacrificed at the indicated time points using CO^2^ inhalation until body functions ceased, followed by cervical dislocation. All experiments were approved by the Local Institutional Animal Care and Research Advisory Committee and authorized by the local government authority.

0.5 mg EdU (Baseclick, Neuried, Germany) was injected intraperitoneally 12 hours before analysis.

For intraperitoneal tamoxifen injections tamoxifen powder (Sigma-Aldrich) was dissolved in corn oil (Sigma-Aldrich) by either incubation at 37°C overnight at 225 rpm or by sonication for 20 minutes in a water bath below 30°C. 1.5 mg of tamoxifen were injected per mouse in 200 µl of tamoxifen.

To generate bone marrow chimeras, mice underwent 2 irradiations of 5Gy each at an interval of 6h. A total of 10^7^ bone marrow cells where intravenously injected within 24h of irradiation. Mice were then given antibiotics in the drinking water (Cotrimoxazol (Ratiopharm)) in a dose of 1.25 mg/ml for 3 weeks after irradiation to prevent opportunistic infections.

For the injection of MC-38 tumors, MC-38 were cultured in DMEM with 10% FCF. Cells were harvested upon reaching 80% confluence after the second passage after thawing. 750,000 cells were subcutaneously injected in the shaved right flank of the recipient mice. Mice were sacrificed when the average tumor volume reached 400 mm^3^ and the tumors and lymph nodes were removed for analysis. One mouse with intrapleural tumor growth was removed from analysis.

### 2. Histology

Proximal small intestine was cut open, rolled, and fixed in 10% formalin for 24-72 hours. After paraffin embedding, 4 µm sections were cut using Microm HM 340E (Thermo Scientific™) and were collected on slides pre-treated with VECTABOND reagent (Biozol, VEC-SP-1800). The slides were dried at room temperature overnight followed by a 1-hour incubation at 56°C.

The tissue sections were deparaffinized by treating the slides with xylene (Otto Fischar GmbH & Co. KG) and in decreasing concentrations of ethanol (VWR) and rehydrated in water. Antigen retrieval was performed by boiling the slides with citrate buffer (10 mM tri-sodium citrate dihydrate (Carl Roth) and 0.05% TweenR 20 (Sigma-Aldrich), pH 6.0) at 75°C for 1 hour. To reduce background noise, the sections were treated with Image-iT FX (ThermoFisher, 136933) for 30 minutes and blocked using PBS and 2% BSA (Carl-Roth, 3737.3) for 1 hour at room temperature to prevent any unspecific binding. To reduce autofluorescence, the sections were treated with TrueBlack (Fisher Scientific, NC1125051). Sections were stained with Ki-67 AF 488 (eBioscience, SolA15) in blocking solution at 4°C overnight and washed several times in PBS and ddH_2_O. Then, the sections were stained in DAPI (Carl-Roth) for 3 minutes at room temperature and once again washed with PBS and ddH_2_O. The slides were embedded in VectaMount Aqueous Mounting Medium (H-5501-60), imaged using a Zeiss microscope (Axio Imager.M2, Carl Zeiss) and analyzed with QuPath (0.5.0).

### 3. Patient samples

Primary colorectal tumors were obtained from routine resections from patients who signed an informed consent approved by the Ethics Committee of the Medical Faculty of the RWTH Aachen University (EK 206/09). Written informed consent was obtained from every patient for research use of the samples. Only patients showing an increase in Tregs in the tumor as compared with unaffected tissue, and having at least 40 Tregs, were retained for analysis.

| patient number | tumor localisation | age | sex |
| --- | --- | --- | --- |
| 1 | proximal<br>(c. ascendens) | 81 | f |
| 2 | proximal<br>(c. ascendens) | 91 | f |
| 3 | proximal<br>(right colic flexure) | 81 | m |
| 4 | proximal<br>(right colic flexure) | 64 | m |
| 5 | distal<br>(recto-sigmoid) | 61 | m |
| 6 | distal<br>(appendix) | 61 | f |
| 7 | distal<br>(left colic flexure) | 88 | m |
| 8 | distal<br>(rectum) | 75 | m |

### 4. Isolation of leucocytes from tissues

Mouse spleen and lymph nodes were meshed up using a syringe stamp on 50µm Nitex in PBS supplemented with 3% fetal calf serum (FCS). Cells were kept on ice. Spleen cells underwent red blood cell lysis with a hypotonic solution (0.17 M NH_4_Cl, 10 mM KHCO_3_, 0.1 mM EDTA). Cell suspensions were washed, resuspended in PBS/FCS, filtered through 50 µm Nitex and counted.

Mouse small intestine (SI) and colon were removed from the peritoneal cavity with as little fat attached as possible. SI was cut into two pieces (proximal SI (pSI) and distal SI (dSI)). Peyer’s patches were removed, the tissue cut open longitudinally and swirled in PBS/FCS until fecal content was washed off. The tissue was then cut into pieces of 0.5-1 cm length and transferred into a 50 ml tube (Falcon) filled with 5-10 ml HBSS (Gibco) /FCS. Intestinal pieces were carefully inverted and then filtered through 50 µm nitex mesh to remove supernatant. Pieces were placed into a fresh tube containing HBSS 2 mM EDTA, shaken vigorously for at least 10 seconds and then incubated at 37°C and 225 rpm for 20 minutes. Tubes were manually shaken every 5-10 minutes. After incubation, tubes were again shaken vigorously, and supernatant removed by filtering through 50 µm nitex. Gut pieces were transferred into room temperature HBSS, shaken strongly, filtered again and then transferred into a fresh tube of HBSS/EDTA. A second round of incubation/shaking in HBSS/EDTA was then performed, followed by a wash in HBSS. Enzymatic digestion was performed at 37°C in RPMI (Gibco) containing 10 % FCS and Collagenase VIII (1 mg/ml; Sigma-Aldrich) for small intestine or Collagenase V (0,85 mg/ml, Sigma-Aldrich), Collagenase D (1.25 mg/ml, Roche), Dispase (1 mg/ml, Gibco) and DNAse (30 mg/ml, Roche) for colon. Tissue was incubated for 20 minutes on 225 rpm at 37°C and shaken manually every 5 minutes. After digestion, the single cell suspensions were washed with PBS/FCS and filtered through a 100 µm cell strainer (Falcon, Corning). Cell suspensions were resuspended in PBS/FCS and counted.

MC-38 tumors were removed from the sacrificed animals, weighed and transferred into 1,5 ml of RPMI/FCS. The tissue was minced using scissors. After mincing, an enzyme cocktail (0.85 mg/ml Collagenase V, 1.25 mg/ml Collagenase D, 1 mg/ml Dispase, 30 µg/ml DNAse) was added. Tissue was incubated at 37°C, 20 minutes shaking at 225 rpm and manually shaken every 4 minutes. Cell suspension was then filtered through a 100 µm cell strainer into a 15 ml tube. To remove tissue debris, a density gradient was performed. For this, a starting Percoll solution (Merck) (100%) was prepared using 45 ml of Percoll solution and 5 ml 10x PBS. A gradient was prepared by resuspending the cells in a solution of RPMI + 40% PBS-Percoll solution and carefully layered on top of a 70% solution of PBS-Percoll with HBSS. After centrifugation, the interface was collected, washed in PBS/FCS and counted.

Human tissue samples were conserved overnight in MACS Tissue storage Solution. 300-600 mg of the tissue were incubated for 30 minutes in HBSS 2mM EDTA, then digested for 20 minutes with Collagenase VIII (1 mg/mL, Sigma-Aldrich), Collagenase D (1.25 mg/mL, Roche) and DNase (30 µg/mL, Roche). Cells were then frozen before analysis.

### 5. Flow cytometry

1 × 10^6^ cells were stained in PBS/FCS with 10% rat serum (Abd Serotec). Cells were then stained with Viability dye as per manufacturer’s instructions. Cells were fixed and permeabilized using the Foxp3/Transcription Factor Staining Buffer Set (ThermoFisher) according to the manufacturer’s protocol. Cells were incubated in 50 µl of intracellular antibody cocktail for 60 minutes. After incubation, cells were washed, resuspended and analyzed in a Cytek Aurora (Cytek Biosciences). Data were analysed using FlowJo Software (BD Biosciences) or OMIQ software (Dotmatics).

For analysis of human Treg in OMIQ, a maximum of 200 cells were concatenated for sample to generate the UMAPs, FlowSOM clustering and heatmaps. Differences in cluster frequencies between samples and volcano plot were generated with EdgeR.

### 6. Isolation of cells and scRNA-seq

To obtain comparable cell numbers, we sorted CD4^+^ CD25^+^ KLRG1^+^ effector tissue Treg from one steady-state and one neoplastic small intestine using a FACSAria II (BD). The choice was made to analyze one individual per group to allow for higher numbers in the clonal analysis, and because of the high penetrance of the phenotype of increased Treg. As controls, we sorted gut CD4^+^ CD25^+^ KLRG1^-^ T cells, which are enriched in other Treg types. Sorted cells were encapsulated into droplets with the ChromiumTM Controller (10x Genomics) and processed following manufacturer’s specifications. Transcripts captured in all the cells encapsulated with a bead were uniquely barcoded using a combination of a 16 bp 10x Barcode and a 10 bp unique molecular identifier (UMI). cDNA libraries were generated using the Chromium™ Next GEM Single Cell 5’ Reagent Kit v1 or v2 (10x Genomics) following the detailed protocol provided by the manufacturer. Libraries were quantified by Quantus (Promega) and quality was checked using TapeStatoin 4200 with High Sensitivity DNA kit (Agilent). Libraries were sequenced with the NextSeq 500 platform (NextSeq 500/550 Mid Output Kit v2.5 (150 cycles), Illumina) with paired-end mode or NovaSeq 6000 platform (SP Reagent Kit v1.5, Illumina) in 50 bp paired-end mode. The hashtag library was demultiplexed using CellRanger software (version 7.1.0). Raw data was demultiplexed with cellranger mkfastq (10x Genomics v7.1.0) to generate fastq files. The fastq files were then analyzed by the cellranger multi with default parameters using refdata-gex-mm10-2020-A as reference.

### 7. scRNA-seq preprocessing and quality control

Single cell analysis was performed using the Scanpy package (v1.9.1, ^48^) in Python (v3.9.4) with all function parameters at default or recommended settings unless described otherwise. Key dependencies for Scanpy included pandas (v1.4.2, DOI: 10.5281/zenodo.3509134), NumPy (v1.20.3,^49^) and Anndata (v0.8.0,^50^). Filtered counts matrices from each run were first subjected to doublet detection using Scrublet (no version number, ^51^) with a manually-adjusted doublet score cut-off threshold determined using scrub.plot_histogram() as described in the example notebook (provided by the developer at https://github.com/swolock/scrublet/blob/master/examples/scrublet_basics.ipynb). Count matrices from all mouse single-cell captures were then merged into a single object and cells were filtered for more than 200 genes and less than 50% mitochondrial reads. Genes expressed in less than three cells were also removed, as well as mitochondrial, ribosomal and immunoglobulin genes.

#### Cell-type annotation and clustering

The merged dataset was then normalised and log transformed as part of the standard Scanpy workflow before being filtered for highly variable genes determined using the command sc.pp.filter_genes_dispersion. The number of counts and percentage mitochondrial gene expression were regressed, and data scaling and principal component analysis were performed. Batch-corrected nearest neighbours were computed using Scanpy’s bbknn function (Polanski et al., Bioinformatics 2020)^52^ and dimensionality reduction via the UMAP function. Leiden clustering was performed using a resolution of 0.3 and cell clusters annotated based on the expression of marker genes identified using the sc.tl.rank_genes_groups function (method= ‘wilcoxon’). Two clusters corresponding to macrophages and B cells were identified, and barcodes corresponding to these cells were flagged for removal. The normalised counts matrix containing only CD4^+^ T cells was then reverted to a pre-normalised state and re-normalised, log-transformed, filtered for highly variable genes and clustered as described above. The cell numbers in the final scRNA-seq analysis are as follows: CD4: 589; CD4 Il7r^+^: 780; Treg cycling: 487; Treg effector Klrg1^+^: 7631; Treg Ikzf2^+^: 476; Treg Sell^+^: 504.

#### T cell repertoire analysis

T cell repertoire analysis was performed using the Scirpy package (v0.10.1, ^53^) and following the standard workflow provided by the developer (accessed at https://scverse.org/scirpy/latest/tutorials/tutorial_3k_tcr.html).

#### Treg scoring

For Treg scoring, the gene lists from ^10,11,14^ were used and are also listed in supplementary table 3. The scoring was done using sc.tl.score_genes() function of Scanpy with default parameters to calculate the average expression of selected genes substrated with the average expression of reference genes.

### 8. TCRa bulk sequencing

UMI-based library preparation is an established technique to reduce PCR and sequencing bias by incorporation of one of > 10^7^ potential unique molecular identifier in each starting cDNA at the first step of TCRa library preparation ^54^. To this purpose, KLRG1^+^ and KLRG1^-^ CD4^+^ CD25^+^ T cells were sorted from small intestinal preparations from 4 neoplastic and 2 steady state mice using FACSAria II (Becton Dickenson. Cells were disrupted by RLT buffer supplemented with 1% β-Mercaptoethanol and RNA was isolated by RNeasy Micro kit (QIAGEN). UMI-based preparation of TCRα and TCRβ full repertoire library was performed using the template switch technique and UMI-adaptor incorporation during the TCR-specific cDNA synthesis (Template Switching RT Enzyme Mix, NEB). cDNA synthesis was followed by two nested PCR amplifications and Illumina adaptor incorporation in a third amplification. PCR products after first and second amplification were purified using QIAquick PCR Purification Kit (QIAGEN). Third PCR products were cleaned up by agarose gel electrophoresis followed by DNA extraction using Monarch® DNA Gel Extraction Kit (NEB). TCR libraries were generated using barcoded primers for multiplex Illumina® sequencing (MiSeq). Libraries were quantified by Quantus (Promega) and libraries pooled for sequencing symmetrically using a MiSeq v3 Reagent Kit.

The data for this study have been deposited in the European Nucleotide Archive (ENA) at EMBL-EBI under accession number PRJEB103021 (https://www.ebi.ac.uk/ena/browser/view/PRJEB103021).

### Materials

Mouse Flow cytometry staining:

|  | Fluorochrome | Clone | Company | Dilution |
| --- | --- | --- | --- | --- |
| CCR8 | Biotinylated | SA214G2 | BioLegend | 1:100 |
| CD137 | PE | 17B5 | BioLegend | 1:200 |
| CD4 | BV605 | RM4-5 | BioLegend | 1:200 |
| CD4 | BV711 | RM4-5 | BioLegend | 1:200 |
| CD4 | BV650 | RM4-5 | BioLegend | 1:200 |
| CD45 | BUV 395 | 30-F11 | BioLegend | 1:100 |
| CD45 | PerCP-Cy5.5 | 30-F11 | BioLegend | 1:200 |
| CD45.1 | BV605 | A20 | BioLegend | 1:200 |
| CD45.1 | APC-Cy7 | A20 | BioLegend | 1:200 |
| CD45.1 | BUV 395 | A20 | BD biosciences | 1:100 |
| CD45.1 | FITC | A20 | BioLegend | 1:200 |
| CD45.2 | BUV 737 | 104 | BD biosciences | 1:200 |
| CD45.2 | BV650 | 104 | BioLegend | 1:200 |
| CD90.2 | BV570 | 30-H12 | BioLegend | 1:200 |
| Foxp3 | eF 450 | FJK-16s | Thermo Fisher | 1:100 |
| Foxp3 | AF488 | FJK-16s | Thermo Fisher | 1:100 |
| FOXP3 | APC | FJK-16s | Thermo Fisher | 1:100 |
| Gata3 | eF660 | TWAJ | Thermo Fisher | 1:100 |
| GATA3 | PE | TWAJ | Thermo Fisher | 1:100 |
| Granzyme A | ef450 | GzA3G8.5 | Invitrogen | 1:100 |
| Helios | PE-Cy5 | 22F6 | Thermo Fisher | 1:100 |
| IL-33Ra (ST2) | BV421 | U29-93 | BD biosciences | 1:50 |
| KI67 | PE-Cy7 | SolA15 | Thermo Fisher | 1:400 |
| KI67 | AF 488 | SolA15 | Thermo Fisher | 1:400 |
| Ki-67 | PerCP-eFluor710 | SolA15 | Thermo Fisher | 1:2000 |
| KLRG1 | BV785 | 2F1/KLRG1 | BioLegend | 1:100 |
| PD-1 | BV711 | 29F.1A12 | BioLegend | 1:200 |
| TCR-β | AF700 | H57-597 | BioLegend | 1:100 |
| TCR-β | APC-Cy7 | H57-597 | BioLegend | 1:200 |
| TCR-β | PE-Cy7 | H57-597 | BioLegend | 1:200 |
| TCR-β | BUV 496 | H57-597 | BD biosciences | 1:100 |
| TCR-β | BV605 | H57-597 | BioLegend | 1:100 |
| MHC II (I-A/I-E) | AF 700 | M5/114.15.2 | BioLegend | 1:200 |
| Streptavidin | BV510 | n/a | BioLegend | 1:200 |
| Streptavidin | PE | n/a | BioLegend | 1:100 |
| Streptavidin | PE-Dazzle594 | n/a | BioLegend | 1:400 |
| Zombie NIR | APC-Cy7/Zombie NIR | n/a | BioLegend | 1:1000 |
| Live/Dead Blue | Live/Dead Blue | n/a | BioLegend | 1:1000 |
| eF780 | APC-Cy7/APC-eF780 | n/a | Thermo Fisher | 1:1000 |

Human Flow cytometry staining:

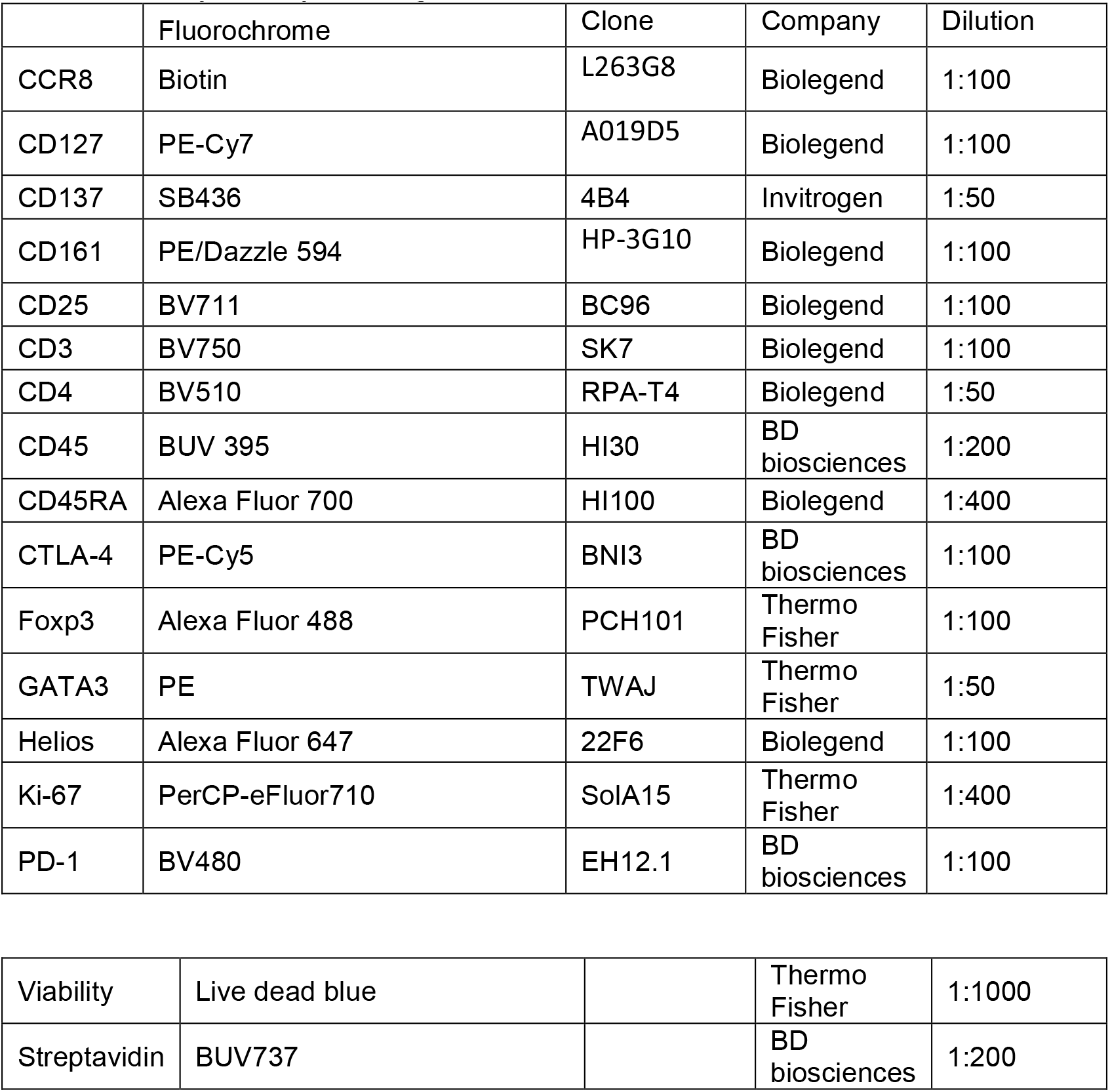

|  | Fluorochrome | Clone | Company | Dilution |
| --- | --- | --- | --- | --- |
| CCR8 | Biotin | L263G8 | Biolegend | 1:100 |
| CD127 | PE-Cy7 | A019D5 | Biolegend | 1:100 |
| CD137 | SB436 | 4B4 | Invitrogen | 1:50 |
| CD161 | PE/Dazzle 594 | HP-3G10 | Biolegend | 1:100 |
| CD25 | BV711 | BC96 | Biolegend | 1:100 |
| CD3 | BV750 | SK7 | Biolegend | 1:100 |
| CD4 | BV510 | RPA-T4 | Biolegend | 1:50 |
| CD45 | BUV 395 | HI30 | BD biosciences | 1:200 |
| CD45RA | Alexa Fluor 700 | HI100 | Biolegend | 1:400 |
| CTLA-4 | PE-Cy5 | BNI3 | BD biosciences | 1:100 |
| Foxp3 | Alexa Fluor 488 | PCH101 | Thermo Fisher | 1:100 |
| GATA3 | PE | TWAJ | Thermo Fisher | 1:50 |
| Helios | Alexa Fluor 647 | 22F6 | Biolegend | 1:100 |
| Ki-67 | PerCP-eFluor710 | SolA15 | Thermo Fisher | 1:400 |
| PD-1 | BV480 | EH12.1 | BD biosciences | 1:100 |
| Viability | Live dead blue |  | Thermo Fisher | 1:1000 |
| Streptavidin | BUV737 |  | BD biosciences | 1:200 |

| Critical commercial assays |  |  |
| --- | --- | --- |
| Chromium Next GEM Single Cell V(D)J Reagent Kits 2 | 10x genomics | Cat#1000265 |
| MinElute PCR Purification Kit | QIAGEN | Cat#28004 |
| MiSeq Reagent Kit v3 | Illumina | Cat#MS-102-3003 |
| NovaSeq 6000 SP reagent Kit v1.5 (100cycles) | Illumina | Cat#20028401 |
| QIAquick® PCR Purification Kit | QIAGEN |  |
| RNeasy Micro Kit | QIAGEN | Cat#74034 |
| Takara ExTaq HS | Clontech | Cat#RR001A |
| Uracyl DNA Glycosylase | New England Biolabs | Cat#M0280S |
| Q5® Hot Start High-Fidelity DNA Polymerase | New England Biolabs | Cat#M0493S |
| Template Switching RT Enzyme Mix | New England Biolabs | Cat#M0466S |
| Monarch®DNA Gel Extraction Kit | New England Biolabs | Cat#T1020L |

### Oligonucleotides

(from ^54^)

#### cDNA synthesis

SmartNNN cDNA amplification forward

AAGCAGUGGTAUCAACGCAGAGUNNNNUNNNNUNNNNUCTT(rG)5

Mus_alfa_synt1

TTTCGGCACATTGATTTG

#### First PCR amplification

Smart 20 forward

CACTCTATCCGACAAGCAGTGGTATCAACGCAG

Mus AV2 rev

GGTGCTGTCCTGAGACCGAG

#### Second PCR amplification

Step_1 (+ Illumina Multiplex Read 1 primer)

ACACTCTTTCCCTACACGACGCTCTTCCGATCT CACTCTATCCGACAAGCAGT

Mus acj (no overlap between nested reverse primer 1. and 2. PCR)

GTGACTGGAGTTCAGACGTGTGCTCTTCCGATCTCAGGTTCTGGGTTCTGGATGT

#### Third PCR amplification

Illumina adapter forward

AATGATACGGCGACCACCGAGATCTACACTCTTTCCCTAC

Illumina adapter reverse

CAAGCAGAAGACGGCATACGAGATCGTGATGTGACTGGAGTTCAGAC

* NNNNN = individual sample barcode

## Abbreviations

Tcon: conventional Foxp3^-^ CD4^+^ T cells
Treg: Foxp3^+^ CD4^+^ regulatory T cells
GATA3 KO: Foxp3 ^YFP-Cre+^ GATA3^fl/fl^ mice lacking GATA3 on all Foxp3^+^ Tregs
FCS: fetal calf serum
SI: small intestine
pSI: proximal small intestine
dSI: distal small intestine

## Notes

https://www.ebi.ac.uk/ena/browser/view/PRJEB103021

### Competing Interest Statement

The authors have declared no competing interest.

