## Supplemental information for "Tumors accumulate expanded GATA3-dependent tissue Tregs"

### Supplemental figures index

Figure S1 Mouse intestinal neoplasia induces the accumulation of Treg resembling human tumor Treg

Figure S2 scRNA-seq of neoplasia-responding Tregs

Figure S3 Clonotype-specific gene expression in effector tissue Treg

Figure S4 Roles of TCR, IL-33R and CD83 in effector tissue Treg response to neoplasia

Figure S5 Role of GATA3 in in effector tissue Treg response to neoplasia

Figure S6 Cell-specific and mono-and biallelic effects of GATA3 in Treg response to tumors

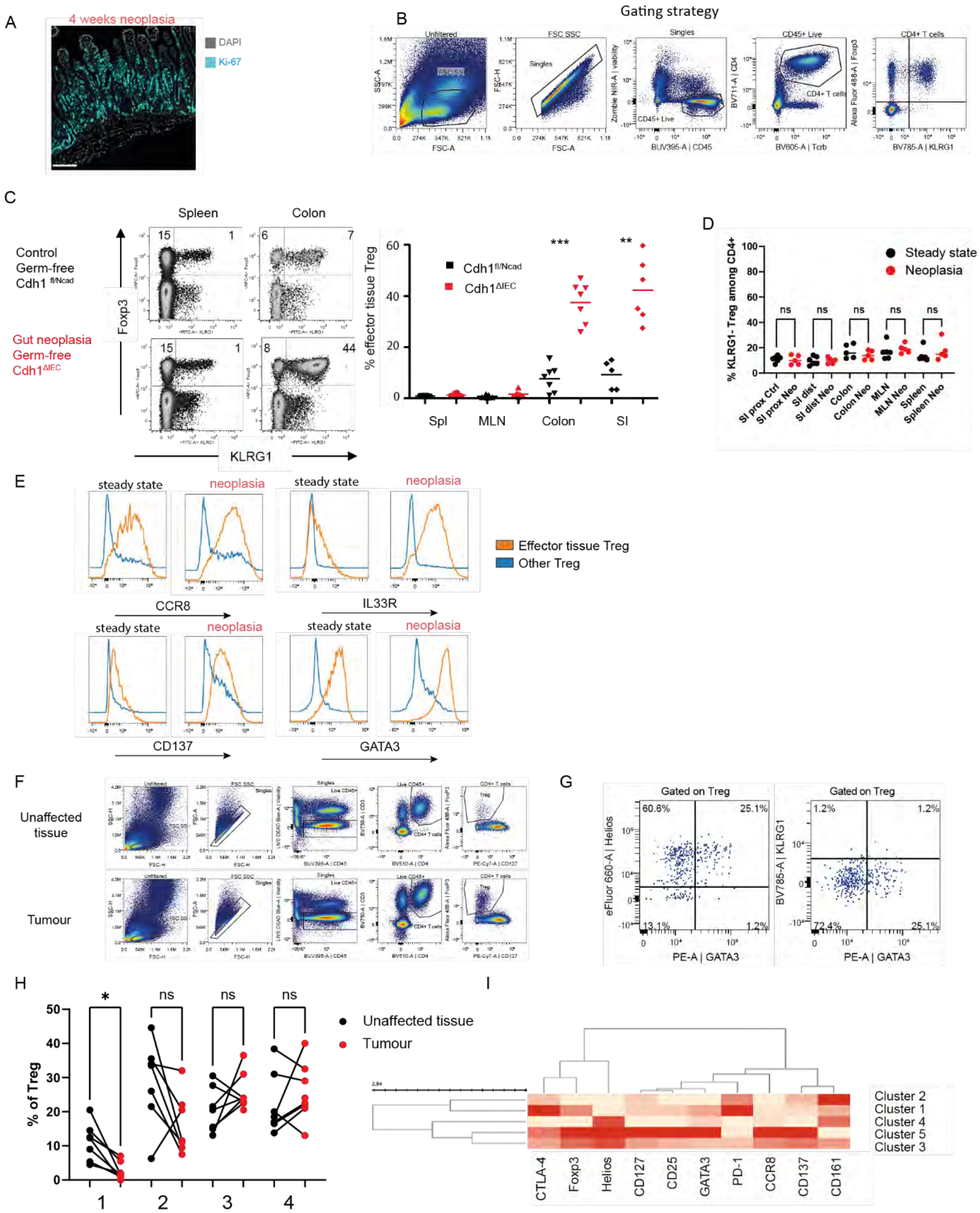


#### Figure S1 Mouse intestinal neoplasia induces the accumulation of Treg resembling human tumor Treg (related to Figure 1)

A: Microscopy image depicting neoplastic changes in the intestinal epithelium 4 weeks after tamoxifen treatment in the *Catnb*$^{+/lox(ex3)}$ Vil Cre-ERT model. Image shows DAPI and Ki67 staining of proximal small intestine as in Fig 1A. Bar: 200 µm.

B: Gating strategy for murine effector tissue and control KLRG1$^{-}$ Tregs. Staining shows proximal small intestine.

C: Representative FACS plots and frequencies of effector tissue Tregs among CD4$^{+}$ T cells in adult germ-free control *Cdh1*$^{fl/Ncad}$ Vil Cre$^{-}$ and cadherin-switched *Cdh1*$^{fl/Ncad}$ Vil Cre$^{+}$ mice (model in [8]). Like the neoplastic model of stabilized β-catenin used in the main results of the article, the *Cdh1*$^{fl/Ncad}$ Vil Cre$^{+}$ model presents increased Wnt signaling in intestinal epithelial cells [8].

D: Frequencies of control non-effector tissue Treg in the indicated organs, 2 weeks after neoplasia induction

E: Representative overlays of IL-33R, CCR8, CD137 and GATA3 expression on effector tissue Tregs (orange, gated as KLRG1$^{+}$ Foxp3$^{+}$) or control Treg (blue, gated as KLRG1$^{-}$ Foxp3$^{+}$) isolated from steady state or neoplastic proximal small intestine 2 weeks after neoplasia induction.

F: Gating strategy of human Treg in unaffected colonic tissue and tumor.

G: Density plot showing GATA3, KLRG1 and Helios expression among Treg cells in human small intestine. Treg were gated as in F. Plot is representative of 2 independent stainings.

H: Frequency of clusters 1-4 from Figure 1L among Treg from human colonic tumors and unaffected tissue. Each pair of dot originates from the same biopsy of one patient.

I: Clustering analysis of human colonic Treg markers in clusters 1-5 from figure 1L.

Each dot in D and E represents an individual mouse, from which all organs shown were analyzed.

A

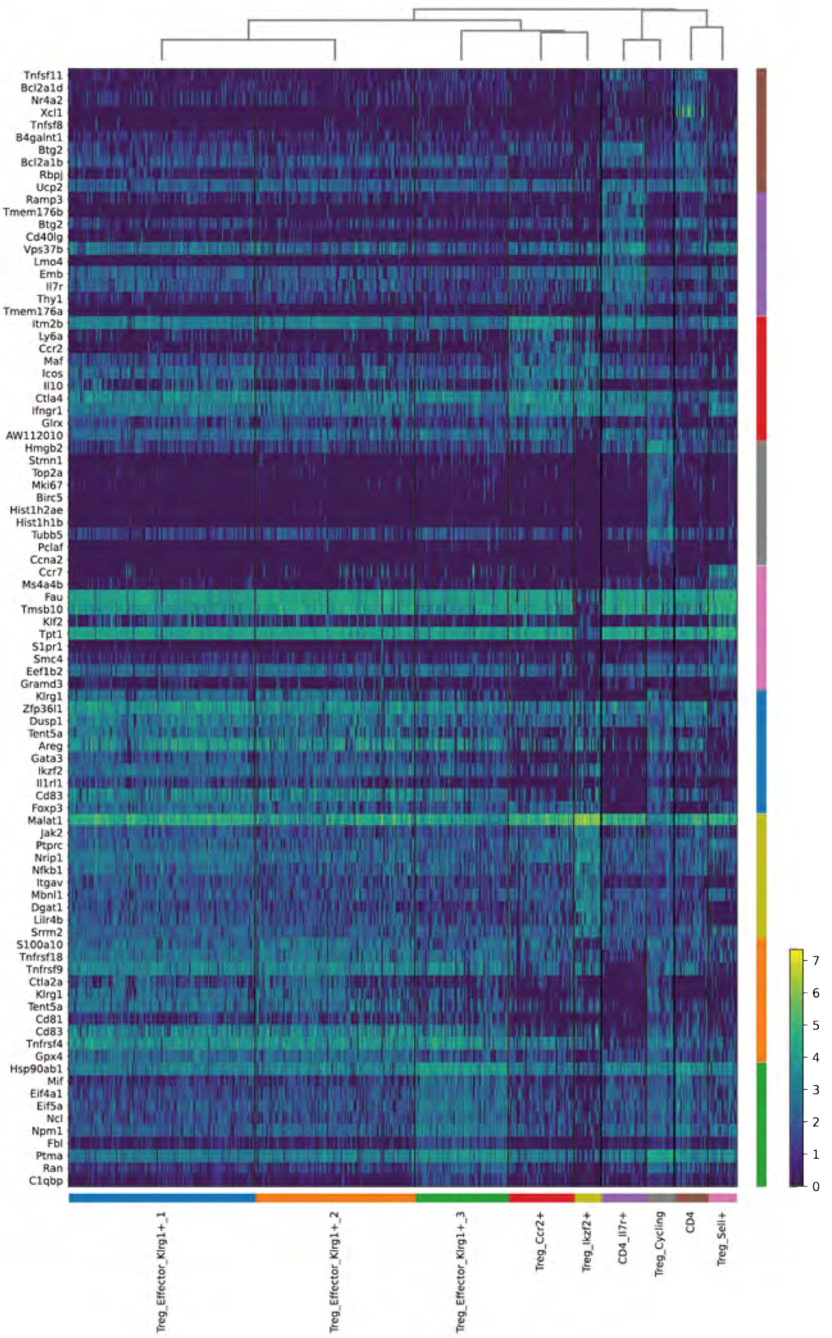


B

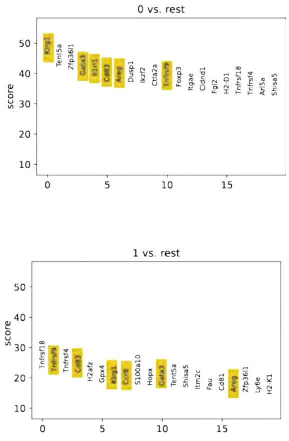


#### Figure S2 scRNA-seq of neoplasia-responding Tregs (related to Figure 2)

A: Expression levels of signature genes for the indicated populations from the clusters in Figure 2.

B: Signature score of the 20 top genes for clusters 0 and 1 from Figure 2.

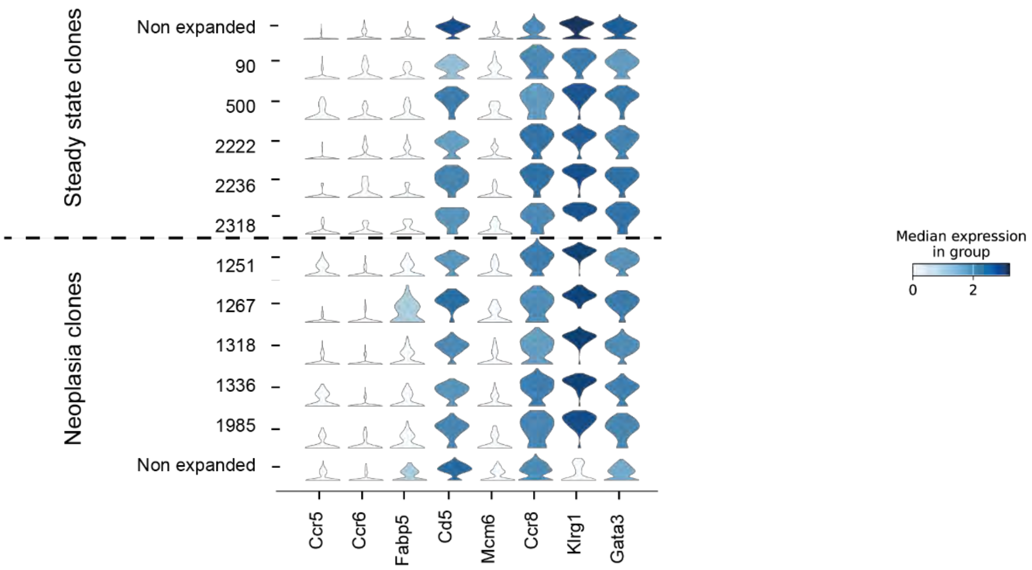


#### Figure S3 Clonotype-specific gene expression in effector tissue Treg (related to Figure 3)

Violin plots showing expression levels of selected genes by the 5 most abundant clonotypes from effector Treg cells isolated from steady-state or neoplastic small intestine, as in Figure 5G. Only cells included in the KLRG1⁺ effector cluster were analyzed. The numbers on the x axis correspond to the clone numbers in Figure 5F.

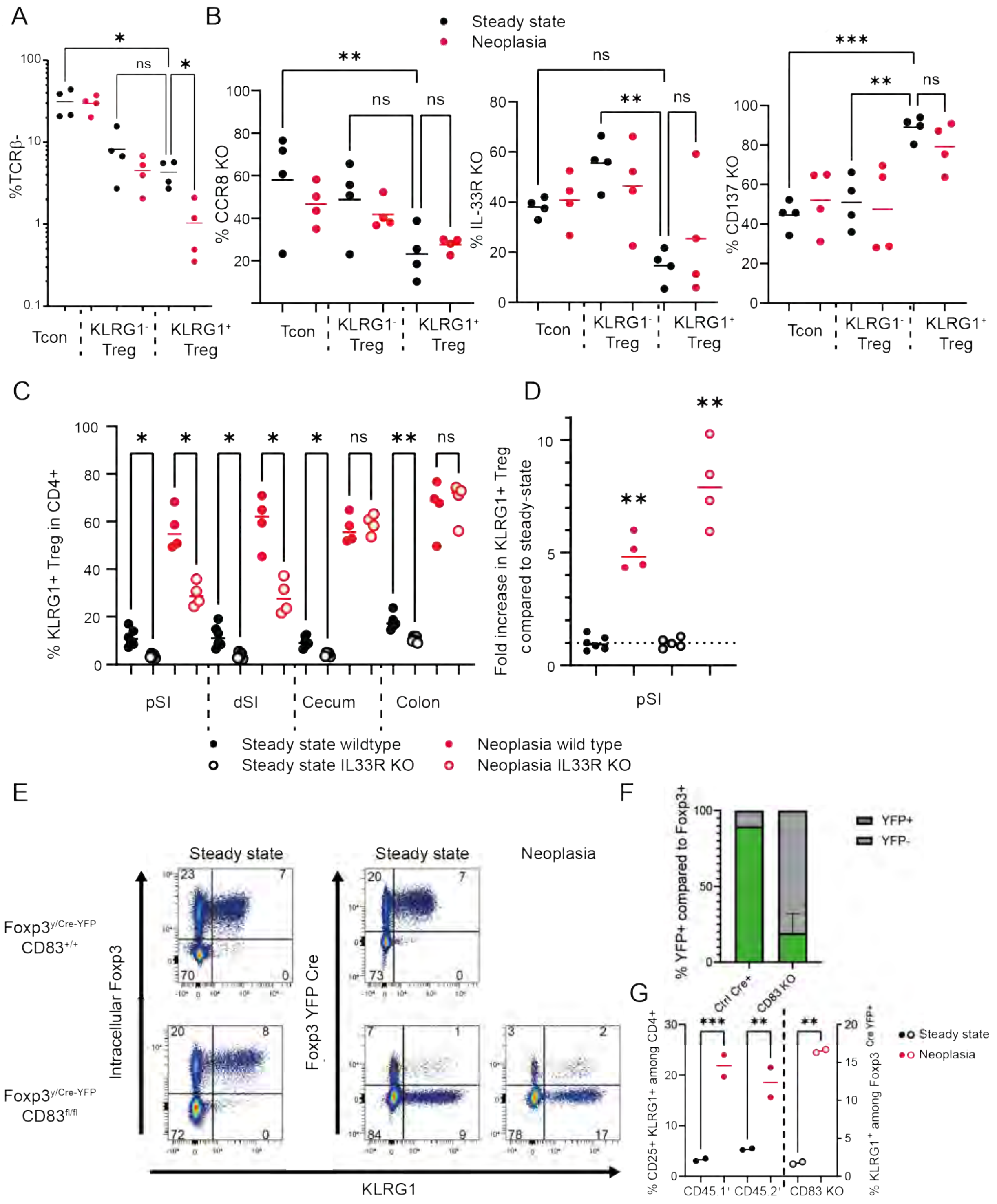


#### Figure S4 Roles of TCR, IL-33R and CD83 in effector tissue Treg response to neoplasia (related to Figure 4)

A: Frequency of TCRβ- cells among CD4⁺ Foxp3⁻ (Tcon), control Treg (CD4⁺ Foxp3⁺ KLRG1⁻) and effector tissue Treg (CD4⁺ Foxp3⁺ KLRG1⁻) from proximal small intestine of CD4$^{Cre/+}$ *Tcra*$^{fl/fl}$ bone marrow chimeras. As a result of TCRβ⁺ KLRG1⁺ Treg expanding during neoplasia, the frequency of TCRβ⁻ among tissue effector Treg is reduced in neoplasia as compared to control intestine.

B: Frequency of CD45.2⁺ KO cells among donor-derived populations in CCR8 KO, IL-33R KO or CD137 KO mixed bone marrow chimeras. The frequencies of KO cells among tissue effector KLRG1⁺ Treg were unaffected by neoplasia in these groups.

C: Frequency of effector tissue Treg among wild-type ($CD45.1^+$) or IL-33R KO ($CD45.2^+$) $CD4^+$ T cells in the indicated organs.

D: Fold increase in $KLRG1^+$ effector tissue Tregs during neoplasia for wild-type and IL-33R KO cells as compared to the mean value of their respective steady states. Statistics assess difference to 1.

E-G: Analysis of CD83 KO mixed bone marrow chimeras into $Catnb^{+/lox(ex3)}$ Vil Cre-ERT recipients.

E: Representative FACS plots showing Foxp3 and KLRG1 expression among $CD45.2^+$ $CD4^+$ T cells from proximal small intestine of CD83 KO ($Foxp3^{YFP\text{-}Cre/y}$ $CD83^{fl/fl}$) or control $Foxp3^{YFP\text{-}Cre/y}$ $CD83^{+/+}$ mixed bone marrow chimeras during steady state or intestinal neoplasia. Intracellular Foxp3: cells were fixed, permeabilized and stained with an anti-Foxp3 antibody. Foxp3 $^{YFP\text{-}Cre}$: Expression of YFP was used as a proxy for Foxp3 expression.

F: $CD4^+$ $YFP^+$ cells calculated as % of $CD4^+$ $Foxp3^+$ cells from steady state mice in E. Ctrl:n= 1, CD83 KO n=2. Frequencies of $YFP^-$ deletion escapees shown in grey.

G: Frequency of effector tissue Treg in steady state and neoplasia among donor $CD4^+$ T cell populations (left) and among $^{YFP\text{-}Cre+}$ CD83 deletors (right).

For all graphs: Black: steady state; red: intestinal neoplasia. Data show one representative experiment out of at least two; each dot represents an individual mouse. Unless stated otherwise, statistics were performed by ANOVA followed by Dunnett's T3 multiple comparisons test, unless stated otherwise. ***, $p<0.001$; **, $p<0.01$; *, $p<0.05$; n.s.: non-significant. For D, one sample t test was performed to assess differences to mean =1.

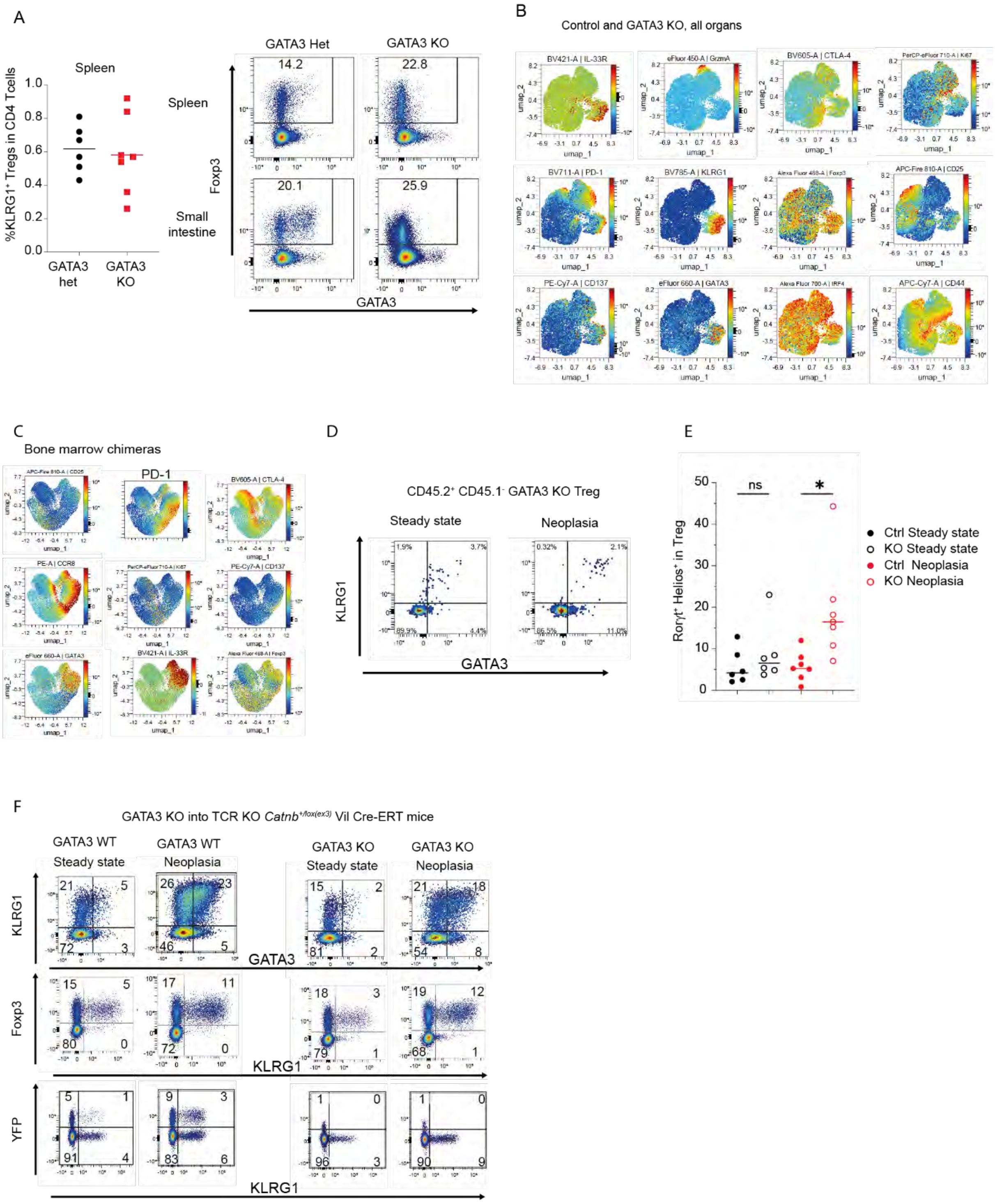


#### Figure S5 Role of GATA3 in in effector tissue Treg response to neoplasia (related to Figure 5)

A: Frequencies of tissue effector Treg in spleen (left) and representative FACS plots of Foxp3 and GATA3 expression showing GATA3 deletion among CD4$^{+}$ T cell from spleen and proximal small intestine (right) of GATA3 Het and KO mice.

B: Heatmap expression of indicated markers on UMAP of concatenated Spleen, MLN, small intestine and colon Treg from 2 wt CD45.1 and 2 GATA3$^{fl/fl}$ Foxp3$^{YFP\text{-}Cre}$ mice. UMAP was performed on IL-33R, GzmA, CTLA-4, PD-1, KLRG1, RORgt, Helios, CD137, CD44 and CD25.

C: Heatmap expression of indicated markers on UMAP of concatenated Spleen, MLN, small intestine and colon Treg from 4 steady-state and 4 neoplastic CD45.1/GATA3 KO mixed bone marrow chimeras was performed on IL-33R, CTLA-4, PD-1, KLRG1, CCR8, RORγt, Helios and CD44.

D: Representative FACS plots showing KLRG1 and GATA3 expression among CD45.2$^{+}$ CD45.1$^{-}$ GATA3 KO CD4$^{+}$ Foxp3$^{+}$ Treg from proximal small intestine of mixed bone marrow chimeras under steady state and neoplastic conditions. Plots are representative of at least 5 individual mice.

E: Frequencies of Helios$^{+}$ RORγt$^{+}$ double positive cells among CD45.1$^{+}$ CD45.2$^{-}$ control (filled circles) or CD45.2$^{+}$ CD45.1$^{-}$ GATA3 KO (empty circles) Tregs from mixed bone marrow chimeras. Tregs were isolated from steady state (black) or neoplastic (red) proximal small intestine.

F: Representative FACS plots showing Foxp3 and KLRG1 expression among CD4$^{+}$ T cells from proximal small intestine of chimeric TCRa KO *Catnb*$^{+/lox(ex3)}$ Vil Cre-ERT or control TCRa KO mice reconstituted with GATA3$^{+/+}$ Foxp3$^{YFP\text{-}Cre/y}$ (GATA3 WT) or GATA3$^{fl/fl}$ Foxp3$^{YFP\text{-}Cre/y}$ (GATA3 KO) bone marrow.

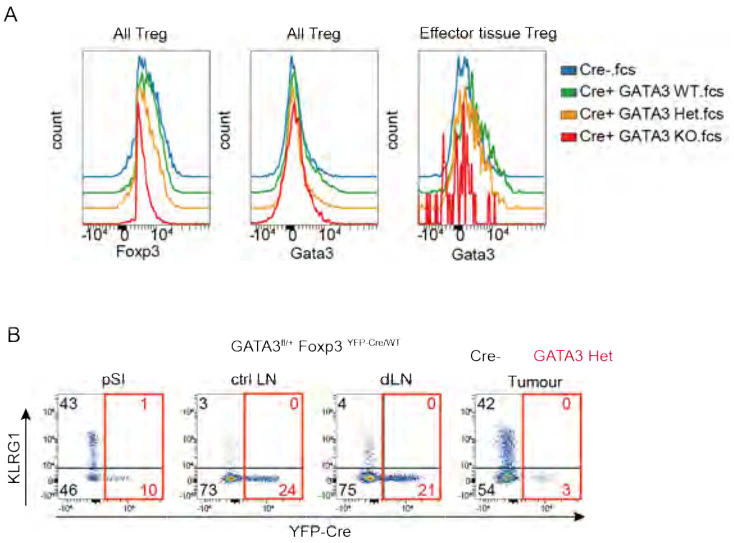


#### Figure S6 Cell-specific and mono-and biallelic effects of GATA3 in Treg response to tumors (related to Figure 6)

A: Representative histograms showing Foxp3 (left) and GATA3 (center) expression in all Treg, and GATA3 expression in effector tissue Treg (gated as Foxp3$^{+}$ KLRG1$^{+}$ CD4$^{+}$ T cells) (right) from proximal small intestine of unmanipulated mice. Red, GATA3$^{fl/fl}$ Foxp3$^{YFP\text{-}Cre\ /YFP\text{-}Cre}$ (GATA3 KO); orange, GATA3$^{fl/+}$ Foxp3$^{Cre\ YFP/YFP\text{-}Cre}$ (GATA3 het); GATA3$^{+/+}$ Foxp3$^{Cre\ YFP/\ YFP\text{-}Cre}$ (GATA3 WT Cre$^{+}$); blue, wild type C57BL/6.

B: Representative FACS plots showing KLRG1 versus *Foxp3$^{YFP\text{-}Cre}$* expression in CD4$^{+}$ CD25$^{+}$ small intestinal T cells from Foxp3$^{YFP\text{-}Cre/+}$ GATA3$^{fl/+}$ females subcutaneously transplanted with MC38 tumor cells. Shown are cells from proximal small intestine, tumor, tumor-draining inguinal lymph node (dLN) and contralateral inguinal lymph node (coLN). Data show one representative mouse out of two analyzed.
