## Supplemental FACS data for "Tumors accumulate expanded GATA3-dependent tissue Tregs"

### TCRa KO

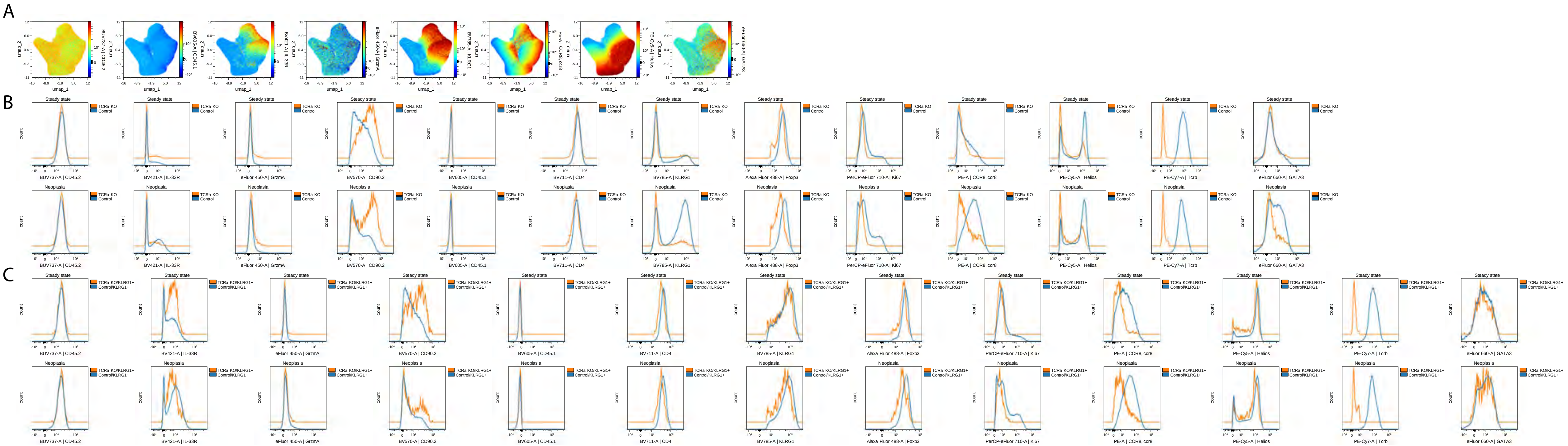


Page 1 FACS analysis of donor-derived Tregs from CD4 CreERT/+ TCRa$^{fl/fl}$ bone marrow chimeras in Catnb$^{+/flox(ex3)}$ Vil Cre ERT mice or Vil Cre ERT- littermates as in figure 4A, 2 weeks after tamoxifen injection. Donor-derived Tregs were gated as CD45.1-, Foxp3+. TCRa KO Treg were gated as TCRβ-, control Treg were gated as TCRβ+ cells. Histograms show concatenated data from 4 control and 4 neoplastic mice.

A: Density UMAP of Treg showing expression of indicated markers.
B: Expression of the indicated markers on total Treg from steady state (top) or neoplastic (bottom) mice
C: Expression of the indicated markers on tissue effector Treg from steady state (top) or neoplastic (bottom) mice. Tissue effector Treg were gated as KLRG1+ cells

### CCR8 KO

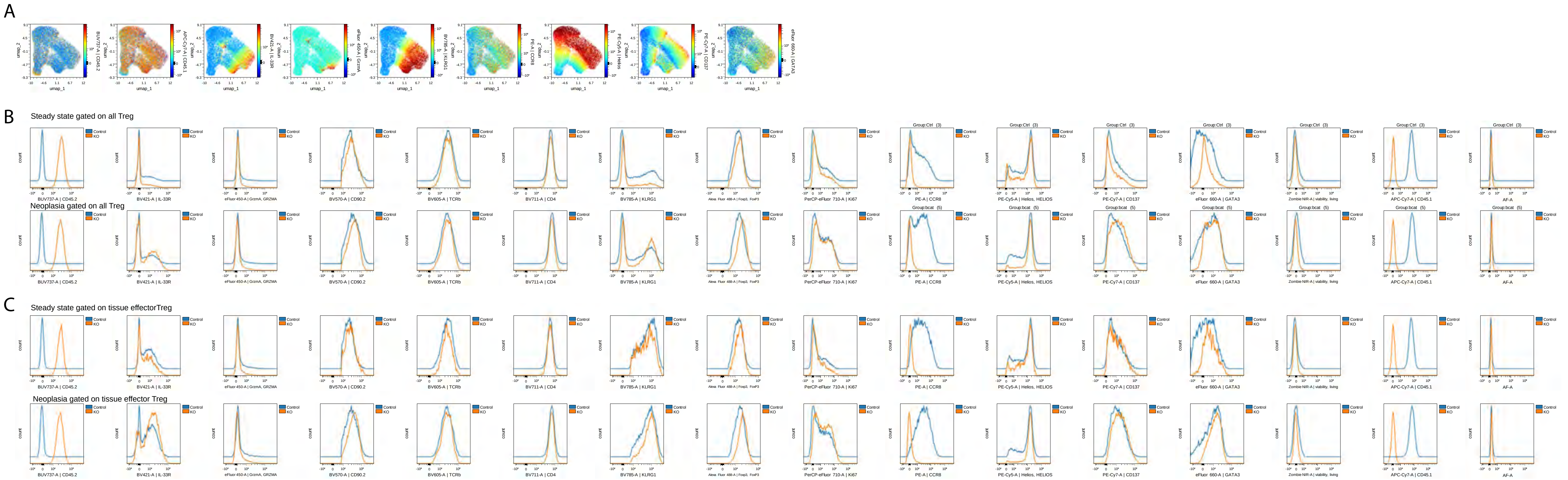


Page 2 Density UMAP of Treg showing expression of indicated markers. UMAP was done on donor-derived Tregs from mixed CCR8 KO/CD45.1+ bone marrow into Catnb$^{+/lox(ex3)}$ Vil Cre-ERT mice or Vil Cre-ERT-negative littermates, 2 weeks after tamoxifen injection as in figure 4E. CCR8 KO donor-derived Tregs were gated as CD45.1$^{-}$ CD45.2$^{+}$ Foxp3$^{+}$. Control donor-derived Tregs were gated as CD45.1+ CD45.2- Foxp3+. Recipient CD45.1+ CD45.2+ cells were excluded from analysis. Histograms show data from 3 control and 5 neoplastic mice.

A: Density UMAP of Treg showing expression of indicated markers.

B: Expression of the indicated markers on total Treg from steady state (top) or neoplastic (bottom) mice

C: Expression of the indicated markers on tissue effector Treg from steady state (top) or neoplastic (bottom) mice. Tissue effector Treg were gated as KLRG1+ cells

# IL33R KO

A

B

C

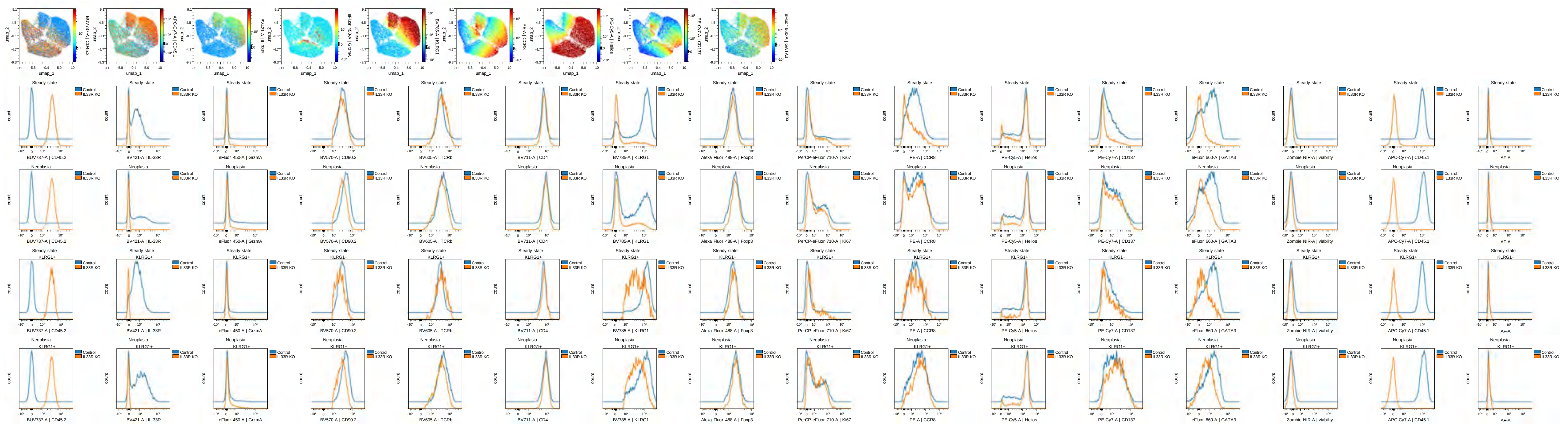


Page 3 FACS analysis of donor-derived Tregs from IL33R KO bone marrow chimeras in Catb$^{flox(ex3)/+}$ Vil Cre ERT mice or Vil Cre ERT- littermates, 2 weeks after tamoxifen injection as in figure 4M.

IL-33R KO donor-derived Tregs were gated as CD45.1-, CD45.2+ Foxp3+. Control donor-derived Tregs were gated as CD45.1+, CD45.2- Foxp3+.

Histograms show concatenated data from 2 mice per population.

A: Density UMAP of Treg showing expression of indicated markers.-

B: Expression of the indicated markers on total Treg from steady state (top) or neoplastic (bottom) mice

C: Expression of the indicated markers on tissue effector Treg from steady state (top) or neoplastic (bottom) mice. Tissue effector Treg were gated as KLRG1+ cells

# CD137 KO

A

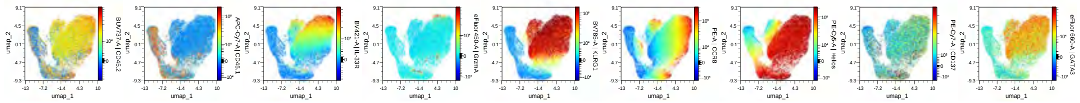

B

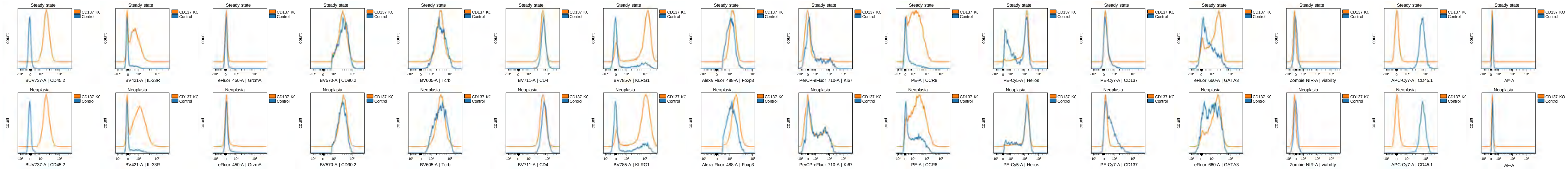

C

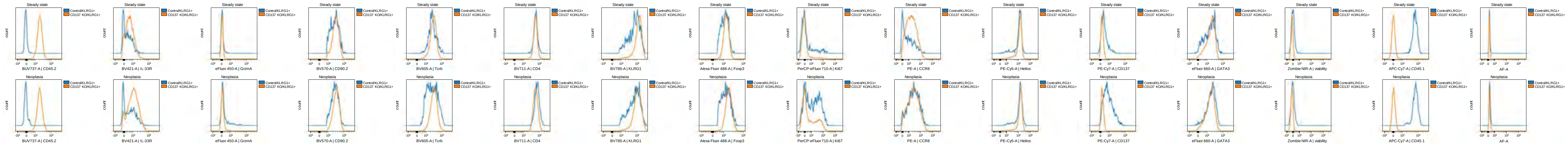

Page 4 Density UMAP of Treg showing expression of indicated markers. UMAP was done on donor-derived Tregs from CD137 KO bone marrow chimeras in Catb$^{flox(ex3)/+}$ Vil Cre ERTmice or Vil Cre ERT- littermates 2 weeks after tamoxifen injection as in figure 4O.
CD137 KO donor-derived Tregs were gated as CD45.1- CD45.2+ Foxp3+. Control donor-derived Tregs were gated as CD45.1+ CD45.2- Foxp3+.
Histograms show concatenated data from 2 mice per group.
A: Density UMAP of Treg showing expression of indicated markers.
B: Expression of the indicated markers on total Treg from steady state (top) or neoplastic (bottom) mice
C: Expression of the indicated markers on tissue effector Treg from steady state (top) or neoplastic (bottom) mice. Tissue effector Treg were gated as KLRG1+ cells

### GATA3 KO

A

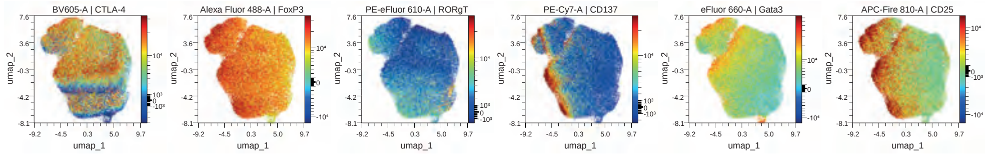


B

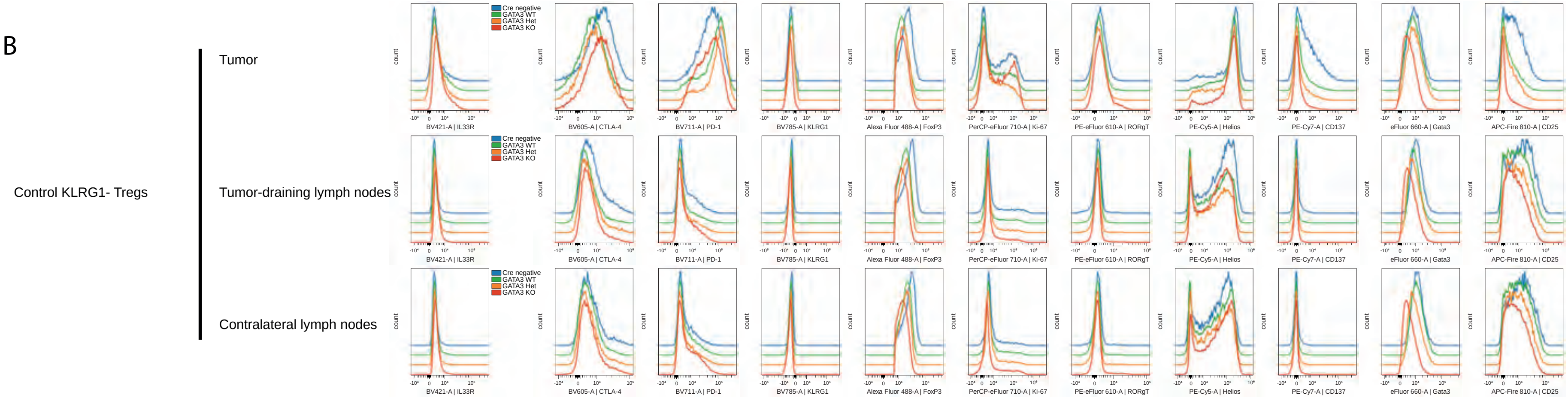


C

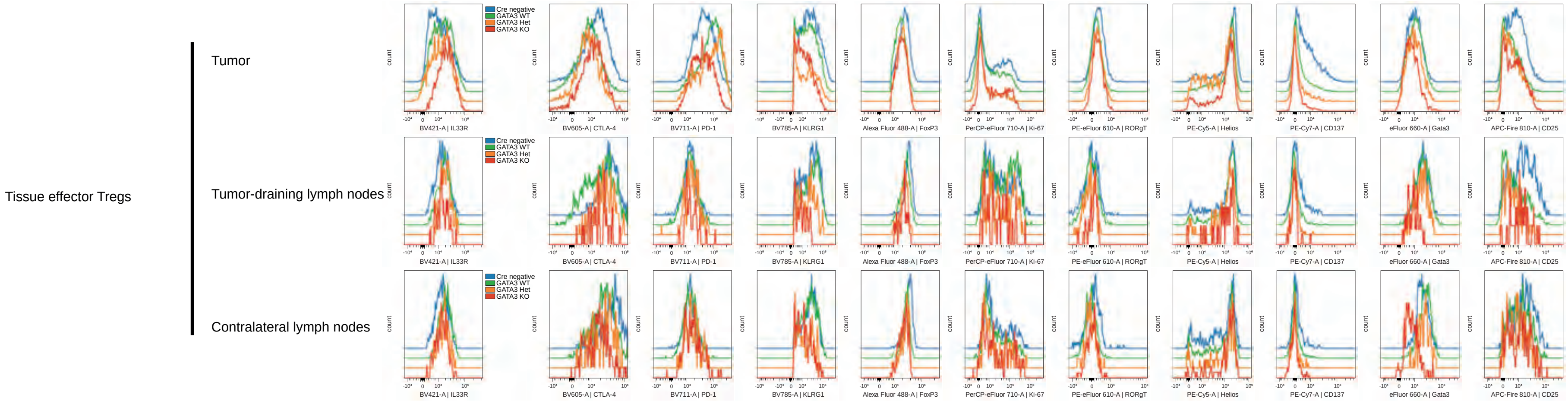


Page 5 Analysis of Tregs from mice with subcutaneous MC-38 tumors
Data are pooled from 2 Cre negative mice, 3 GATA3 WT mice, 2 GATA3 Het mice and 3 GATA3 KO mice.
A: Density UMAP of tumor Treg showing expression of indicated markers as in Figure 6I.

B: Expression of the indicated markers on control KLRG1$^{-}$ Treg from indicated sites.

C: Expression of the indicated markers on tissue effector Treg from indicated sites
